# Neural Dynamics of Multiattribute Decision Making Under Choice Overload

**DOI:** 10.1101/2025.11.18.689103

**Authors:** Jacob M. Stanley, Douglas H. Wedell

## Abstract

The present study examines the neural mechanisms of value-based multiattribute decision making through the lens of choice overload during consumer choice. Two behavioral experiments established that choice sets of a moderate size with nine options were perceived as most optimal compared to choice sets with three and 24 options, and options chosen from the largest set size were judged to be the least satisfying. A functional MRI study then examined how the brain responds to choice sets varying in size and complexity by manipulating the number of options presented as well as the presence of asymmetrically dominated decoy alternatives during a simulated online shopping task. Results showed that the dorsolateral prefrontal cortex (DLPFC) activity followed an inverse U-shape as a function of choice set size, peaking for moderately sized choice sets. The anterior cingulate cortex (ACC) exhibited a linear trend with choice set size. Trials containing decoy options elicited greater activation of the anterior insula (AIns) and DLPFC. Computational modeling of choice behavior revealed a greater tendency to utilize a simplifying lexicographic decision strategy as decision difficulty increased, and individual differences in decision strategies were reflected in activity of the ACC and AIns. These findings advance understanding of how the brain integrates effort, control, and strategy during complex value-based decisions.

---

Choice overload is a mental state that occurs when the complexity of a decision problem exceeds the cognitive resources of the decision maker (Chernev et al., 2015). This has commonly been referred to as the paradox of choice in which having more options is not always better (Schwartz, 2004). Choosing from large assortments has been shown to produce detrimental outcomes including decreased choice satisfaction (Chernev et al., 2015), increased stress and anxiety (Nagar & Gandotra, 2016; Saltsman, 2019; Schwartz, 2000; Schwartz, 2004), and an increased likelihood of deferring choice altogether (Iyengar & Lepper, 2000; Iyengar et al., 2004). Recent technological advancements have vastly expanded access to diverse and abundant markets. As a result, scenarios of choice overload have spread to new domains and become more prevalent than ever before (Schwartz, 2004). Company management (Zeike et al., 2019), insurance plan selection (Peters et al., 2007), public policy (Czech, 2016), prosocial behavior (Herzenstein et al., 2020) and consumer behavior (Chernev et al., 2015) are all affected by the expansion of choice. Although behavioral outcomes of choice overload are well studied, there is a paucity of studies on how the human brain responds to decisions that elicit choice overload. The present study utilizes functional magnetic resonance imaging (fMRI) to examine the neural processing of decision-making during scenarios relating to choice overload.

Thus far, a single experiment has used fMRI to identify brain regions in humans that are involved in experiences of choice overload by comparing neural activity during decisions among choice sets of different sizes (Reutskaja et al., 2018). Their research showed that choice sets of moderate size elicit greater activity in the ACC, the DLPFC, and the dorsal striatum (bilateral caudate nucleus (CN) and putamen) compared to choice sets of smaller and larger sizes, resulting in an inverse U-shape pattern of activation as a function of choice set size. This single peak function of neural activity in the ACC, DLPFC, and dorsal striatum observed by Reutskaja et al. (2018) is hypothesized to be a neural expression of the Theory of Preference (Coombs & Avrunin, 1977). Together, peak activity in these areas provides evidence for maximal engagement with the decision process and an optimal point for accruing the benefits of a larger choice set while avoiding the detriments of choice overload.

It is important to note that the observation of a quadratic activity profile in the ACC, striatum, and DLPFC reported by Reutskaja et al. (2018) was for choices in the domain of aesthetic preference for perceptual images of landscapes. While these landscape images are useful stimuli for capturing subjective visual preferences, the present study examines if similar mechanisms can be used to characterize the more commonly studied multialternative, multiattribute choice paradigm, in which multiple options are described on multiple attributes for deliberation in domains such as consumer choice. Multialternative, multiattribute choices are designed to be representative of the decisions made by individuals in everyday life, and a large body of literature supports their utility for examining contextual choice (Huber & Puto, 1983; Simonson, 1989; Simonson & Tversky, 1992; Trueblood et al., 2013; Wedell, 1991). Options in these choice sets often have well-defined numerical attributes that facilitate a utilitarian approach to the decision process. Perceptual information, like that conveyed by landscape images, has been shown to be integrated into decision processes differently than numerical representations and produce distinctly different contextual effects (Frederick, 2014). This difference suggests that numerical and perceptual information may link to different neural computational processes. For example, perceptual preference may be more automatically driven whereas multiattribute choice may be more conscious and resource intensive (Kahneman, 2011). Therefore, an examination of the neural processing for the numeric attributes in a multialternative, multiattribute choice is necessary for a more complete understanding of the mechanisms driving experiences of choice overload. Observation of a single peak function of neural activation on assortment size with numerical attributes in a consumer choice paradigm would provide evidence for a domain general role of the ACC, DLPFC, and dorsal striatum in the integration of costs and benefits of decision processes and in experiences of choice overload. A failure to observe a single peak function would demonstrate a need for a more nuanced theory for the neural representations of choice overload.

A meta-analysis of choice overload identified four moderating factors of the effect of the number of options on experiences of choice overload (Chernev et al., 2015). According to this model, choice set complexity, decision task difficulty, preference uncertainty, and decision goal all affect the impact of assortment size on choice overload. The moderating relationship of choice set complexity has been leveraged to accurately identify brain regions linked to experiences of choice overload (Chernev et al., 2015; Reutskaja et al., 2018). Choice complexity can be lowered by adding dominating alternatives into a choice set (Chernev et al., 2015). Dominating alternatives are options that are strictly superior to other options (e.g., of the same quality but at a cheaper price) and they can lower decision complexity by reducing the processing costs required by the decision problem (Chernev, 2006). In the case of landscape images used by Reutskaja et al. (2018), dominating alternatives, or “clear favorites,” were defined as those that were most preferred for each participant in a prior judgement task. Trials that included a dominating landscape image elicited significantly greater activity of the ACC, dorsal striatum, and DLPFC than trials that did not (Reutskaja et al., 2018). Lowering the choice set complexity by adding a dominating option reduced the cost of choosing, which increased the perceived subjective value of the choice set. This was reflected by an increase in activity of the key brain areas showing quadratic activity relationships as a function of choice set size, providing further evidence of their linkage to experiences of choice overload.

Similarly, research on dominating alternatives in multialternative, multiattribute choice have observed greater activity in these same brain areas for choice sets with dominating alternatives than for choice sets without them (Busemeyer, 2019; Hedgcock & Rao, 2009). However, thus far, multialternative, multiattribute stimuli have not been used to study the neural correlates of choice overload, partly due to the difficulty in controlling the variability generated by adding alternatives to choice sets (i.e., more alternatives create more possible configurations of attribute values). For this reason, almost all research on multialternative, multiattribute decision-making has been conducted on choice sets with only three alternatives. To address this limitation, we developed a novel method for studying context effects in a multialternative, multiattribute decision-making task for choice sets with more than three alternatives (Stanley & Wedell, 2024). Our method adds alternatives to the choice set in multiples of three so that, as the choice set increases in size, the proportion of dominating alternatives remains the same and thus preserves comparability across choice sets. This method allows the present study to simultaneously fill two gaps: determining the generalizability of neural mechanisms of choice overload across different types of stimuli (perceptual vs numerical) and the study of decision-making strategies across large choice sets for multialternative, multiattribute choice.

The present study joins research on choice overload and multialternative, multiattribute decision-making, using a novel method developed by our lab for studying asymmetrically dominating alternatives within different sized choice sets (Stanley & Wedell, 2024). We use fMRI to measure brain activity while participants choose consumer products from small, moderate, and large assortment sizes which include dominating alternatives on half of all trials. By employing a combination of fMRI, behavioral measures, and computational modeling across three experiments, this study provides a comprehensive analysis of the neural mechanisms behind choice overload. This integrative approach advances our understanding of the decision-making process at the neural level and offers valuable insights for fields such as behavioral economics and consumer psychology. These insights are essential for designing environments that help individuals make better choices, thus improving consumer well-being, as well as facilitating targeted treatment and interventions for individuals with impairments in brain regions implicated in value-based decision making.

## Behavioral Experiments

We conducted two behavioral experiments in order to establish a foundation for our neuroimaging study on the neural correlates of choice overload in multiattribute choice. The goal of the behavioral experiments was to gain an understanding of the factors that influence the perception of multiattribute choice sets. Across the two behavioral studies, we manipulated the number of options, the presence of decoys, time constraints, and the contextual relationships between options while collecting choice preference data and judgements about the characteristics of the decision-making process.

### Behavioral Experiment 1

Behavioral Experiment 1 aimed to determine, for the grocery stimuli used across all experiments in the present study, how many options are perceived as too few, too many, and optimal. This study manipulated the number of alternatives in eight conditions: three, six, nine, 12, 15, 18, 21, and 24 options. We hypothesized that three options would be rated as too few options and that 24 options would be rated as too many. We further hypothesized that 12 options would be perceived as optimal based on results from Reutskaja et al. (2018) that the choice sets containing 12 landscape images were rated as most ideal. Additionally, Behavioral Experiment 1 sought to understand how the contextual relationship between options in a choice set would affect judgements of the choice set and decision process. To investigate this question, we manipulated the attribute values of the available options in three configurations: one equi-preference contour, three parallel equi-preference contours, and randomized values.

### Methods

#### Participants

A total of 68 participants (age range, 18 – 53 years; mean age, 20.6 years; 5 males, 1 prefer not to say) from the University of South Carolina Sona participant pool were surveyed online using the Qualtrics survey software. Participants were required to complete the experiment in one sitting, and they were not allowed to use a smartphone internet browser. Of the 68 participants surveyed, 26 participants were excluded based upon the Qualtrics durations indicating they had spent more than 60 minutes (indicating that they were distracted during the experiment or were not completing it in one sitting). Thus, a total of 42 participants’ data were used in the final analysis. Participants earned course credit for participating.

#### Procedure

Participants signed up for the experiment on the University of South Carolina Sona participant recruitment website and were given a link to the Qualtrics survey. To begin the survey, they accepted an informed consent document and then were given written instructions on their task. The experiment had participants simulate shopping online for grocery items. Participants were told that they were shopping for groceries online and to select the alternative with the most attractive price and quality rating combination. Each trial presented a screen with a generic picture of the grocery item and either three, six, nine, 12, 15, 18, 21, or 24 alternatives, each with a different price and quality rating. Figure 1 illustrates an example trial with nine options and three equi-preference contours. The arrangement of the available alternatives on screen was randomized on each trial. Participants used their mouse cursor to select their preferred alternative and then clicked an arrow button to continue to the next trial. Participants were required to spend a minimum of 10 seconds and a maximum of 15 seconds on each choice. If the participant did not submit their answer by the 15 second time limit, the experiment progressed, and their answer was submitted as their most recently clicked option. After each choice, participants were asked “How difficult was it to make your decision?” (1-10 scale from Not at all difficult to Extremely difficult) and “Did you feel you had the right number of options to choose from?” (1-9 scale where 1 indicated too few options, 5 indicated just the right amount of options, and 9 indicated too many options).

**Figure 1.**
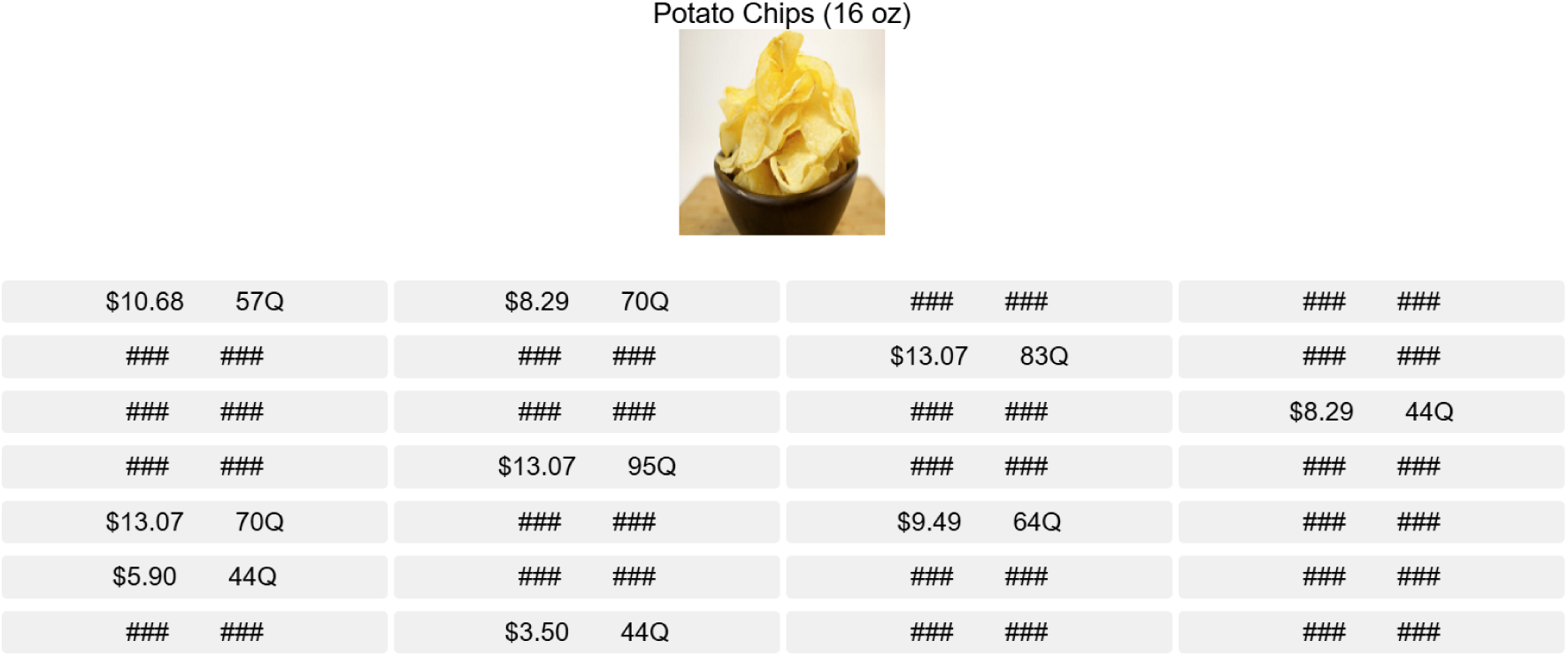
Display of nine options in Behavioral Experiment 1

#### Materials and Designs

Stimuli were based on those used by Wedell et al. (2022). Each option has two attributes, price and quality. Prices are designed to reflect the range of real-world prices for the item. Quality ratings fall on a 100 point scale and represent things characteristics such as brand and spoilage of the item. More information about how the stimuli attribute values were constructed can be found in the supplemental materials. Figure 2 provides a visual representation of the design values used for one contour, three contours, and randomized choice sets.

**Figure 2.**
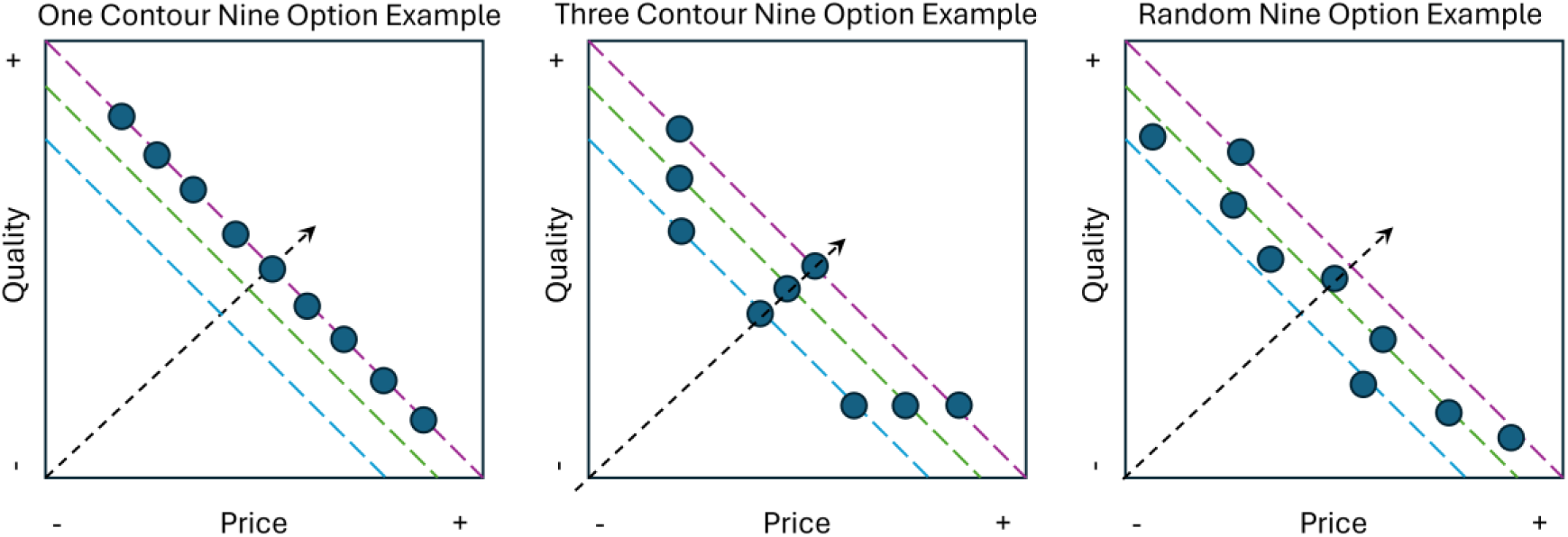
Nine option example of design values for attributes in Behavioral Experiment 1

### Results

To test the effect of including dominance relationships between alternatives on perceived difficulty, we conducted a two-way Number x Contour repeated measures ANOVA on ratings of difficulty for choices made from one contour and choices made from three contours at each level of option number. Note, two participants were excluded from this analysis as they failed to provide a response to some questions. A significant main effect of Contour revealed that choices made from 1 contour were rated significantly more difficult than those made from 3 contours, *F*(1, 39) = 13.77, *p* < .001, η_p_^2^ = 0.26. Panel ‘a’ in Figure 3 shows that the main effect of including dominated alternatives effectively decreased the difficulty of the task. A significant main effect of Number confirmed that the number of options significantly influenced ratings of difficulty, *F*(7, 273) = 23.23, *p* < .001, η_p_^2^ = 0.37. The two-way interaction of Number and Contour was not significant.

**Figure 3.**
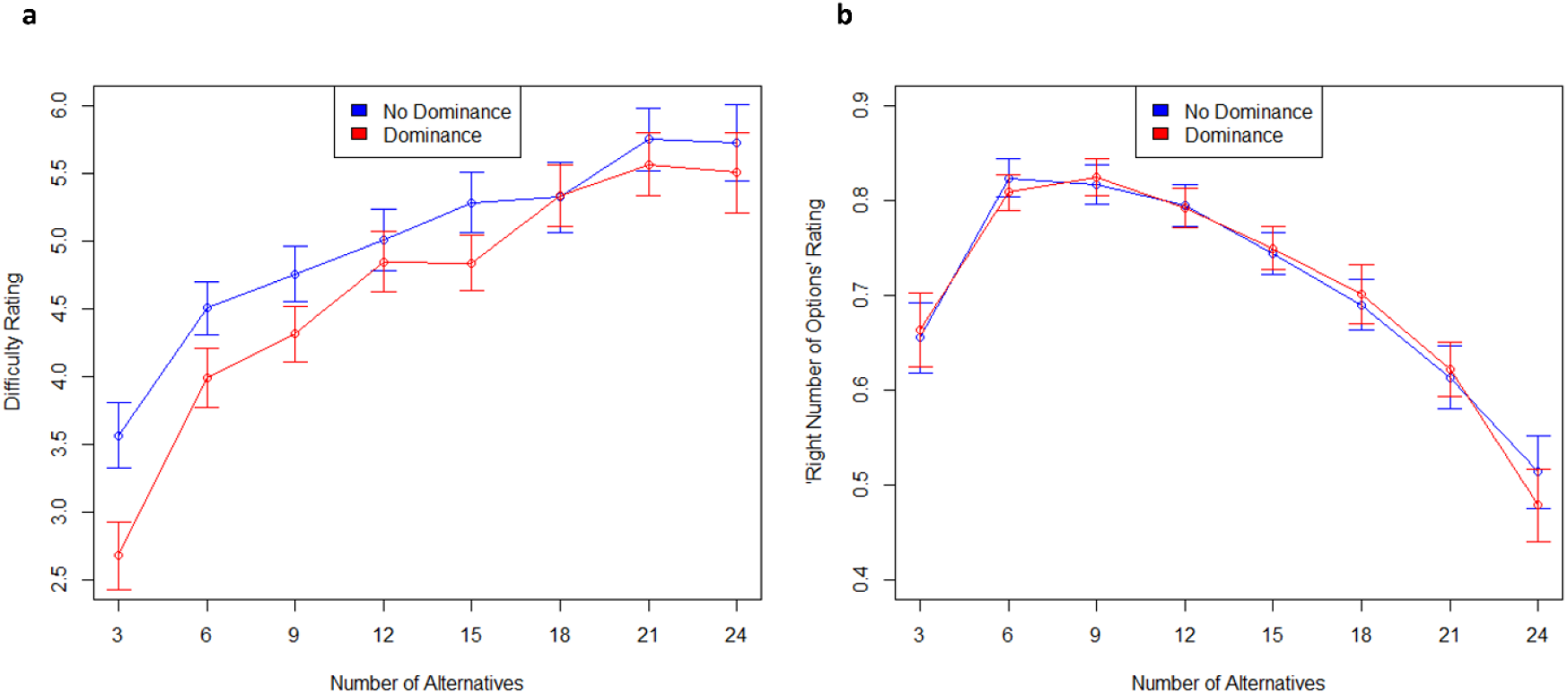
Judgement Ratings for Behavioral Experiment 1. Difficulty ratings show the average perceived choice set difficulty as a function of choice set size are displayed in panel ‘a’. The blue lines show ratings for trials without dominating alternatives. The red lines show ratings for trials with dominating alternatives. Average ‘Right number of options’ ratings as a function of choice set size are displayed in panel ‘b’. A rating of 1 indicates being closest to the right number of options, and a rating of 0 indicates being the furthest. Error bars represent standard errors of the mean ratings.

Panel ‘b’ in Figure 3 plots the degree to which assortment size was perceived as the ‘right number of options’ for our stimuli. For interpretability, we convert the 1-9 scale as

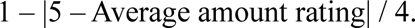

This index displays ratings on a 0 to 1 scale where 1 indicates being perceived as the most optimal and 0 indicates being perceived as the least optimal number of alternatives. We find that six, nine, and 12 alternatives are all perceived to be close to optimal and three and 24 alternatives are clearly perceived to be nonoptimal. In our neuroimaging study we used choice sets of three, nine and 24 alternatives for comparability with Stanley and Wedell (2024). Ratings given to three, nine, and 24 alternative choice sets are also close in value to those given to small, medium, and large choice sets for the stimuli used by Reutskaja et al. (2018).

### Behavioral Experiment 2

Behavioral Experiment 2 sought to examine the effect of time constraints on judgements of small (three), medium (nine), and large (24) choice sets. It also examined the influence of decoys on choice preferences at each level of option number.

### Methods

#### Participants

A total of 53 (M_age_ = 19.9, SD_age_ = 2.2) participants from the University of South Carolina Sona participant pool were surveyed online using the Qualtrics survey software. Participants were required to complete the experiment in one sitting, and they were not allowed to use a smartphone internet browser. Participants earned course credit for participating. No participants were dropped for failing to meet survey requirements so that analyses were based on the full set of 53 participants.

#### Procedure

Participants completed 72 trials of a 3 (number of options: three, nine, 24) x 2 (decoys, no decoys) x 2 (timing: 15s, self-paced) mixed factorial design. Trials presented choice sets with either three, nine, or 24 options within-subjects. Choice sets either had all options trade-off equally on price and quality (no decoy) or contained decoy options that were dominated on one attribute and tied on the other (decoy) within-subjects. Nested within the decoy factor, the trials varied which options were dominating, either high-quality, expensive options (group A) or low-quality, cheap options (group B). Participants either made all decisions within a 15 second timeframe (timed) or at their own pace with no time limit (self-paced), between subjects. After every trial, participants answered the same three trial judgement questions used in Experiment 1. After these choices trial judgments participants filled out a measure of grit (Grit-S) and a measure of cognitive reflection (Cognitive Reflection Test).

#### Materials and Design

Stimuli were based on those used by Wedell et al. (2022). Price and quality attributes for the core alternatives within each choice set were generated in the same way that attributes were generated for options in the single contour condition for behavioral experiment 1, with 𝑁 equal to the number of core alternatives in the choice set. A core alternative is any option that is not a decoy. Attribute values for decoy options were created as follows. Let 𝐷_𝑃_and 𝐷_𝑄_ denote the price and quality of the decoy, let 𝑇_𝑃_ and 𝑇_𝑄_ denote the price and quality of the target, and let 𝑅_𝑃_and 𝑅_𝑄_ denote the price and quality range for the real grocery item. Price and quality were generated for decoys depending upon the attributes of the target alternative. For an attraction decoy favoring A, attributes were calculated as 𝐷_𝑃_ = 𝑇_𝑃_ + (0.20 ∗ 𝑅_𝑃_) for price and 𝐷_𝑄_ = 𝑇_𝑄_ for quality. For an attraction decoy favoring B, attributes were calculated as 𝐷_𝑃_ = 𝑇_𝑃_for price and 𝐷_𝑄_ = 𝑇_𝑄_ − (0.20 ∗ 𝑅_𝑄_) for quality. Figure 4 provides a visual representation of the spatial relationship between options in each experimental condition.

**Figure 4.**
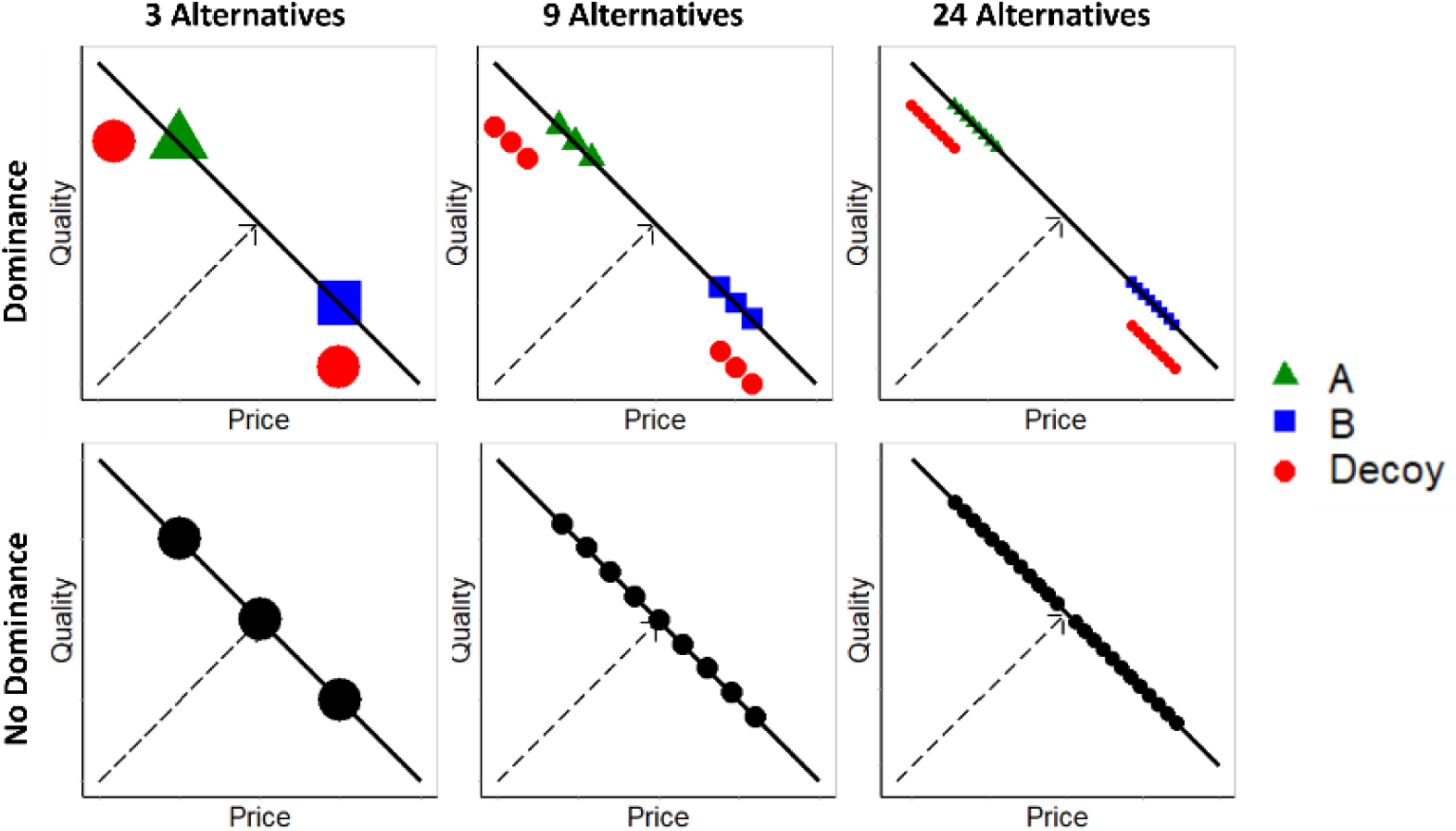
Stimulus Attribute Designs. Stimulus attribute designs are displayed for each experimental condition. Note that the price x-axis becomes better (cheaper) from left to right. In the upper row, both decoy group types (targeting A and targeting B) are shown in each panel where only one is included on any given trial.

### Results

To examine the effect of the decoys on choice preferences, Decoy Effect scores were calculated in the same manner as for Stanley and Wedell (2024) which is as follows:

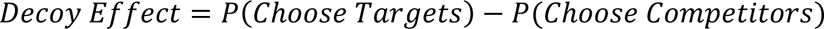

These decoy effect scores are calculated for each participant and for each level of the number of options factor. Decoy Effects can range from −1 to 1. A positive score indicates that a participant is showing the attraction effect, and a negative score indicates a reversal of the expected attraction effect, referred to as the repulsion effect. A zero score indicates that the decoys had no influence on the relative choice proportions of the available alternatives.

A single sample t-test was performed for each experimental condition on Decoy Effect scores with null hypothesis µ = 0. Significant attraction effects were found in choice sets with three, *t*(52) = 5.51, *p* < 0.001, *d* = 0.76, and nine, *t*(52) = 3.62, *p* < 0.001, *d* = 0.40, options but not for those with 24 options. A one-way ANOVA revealed a significant main effect of the number of options on the decoy effect, *F*(1, 51) = 20.32, *p* < 0.001, η^2^_p_ = 0.28 indicated that choice preferences are shifted less with greater numbers of options. Figure 5 displays the choice proportion data for each experimental condition.

**Figure 5.**
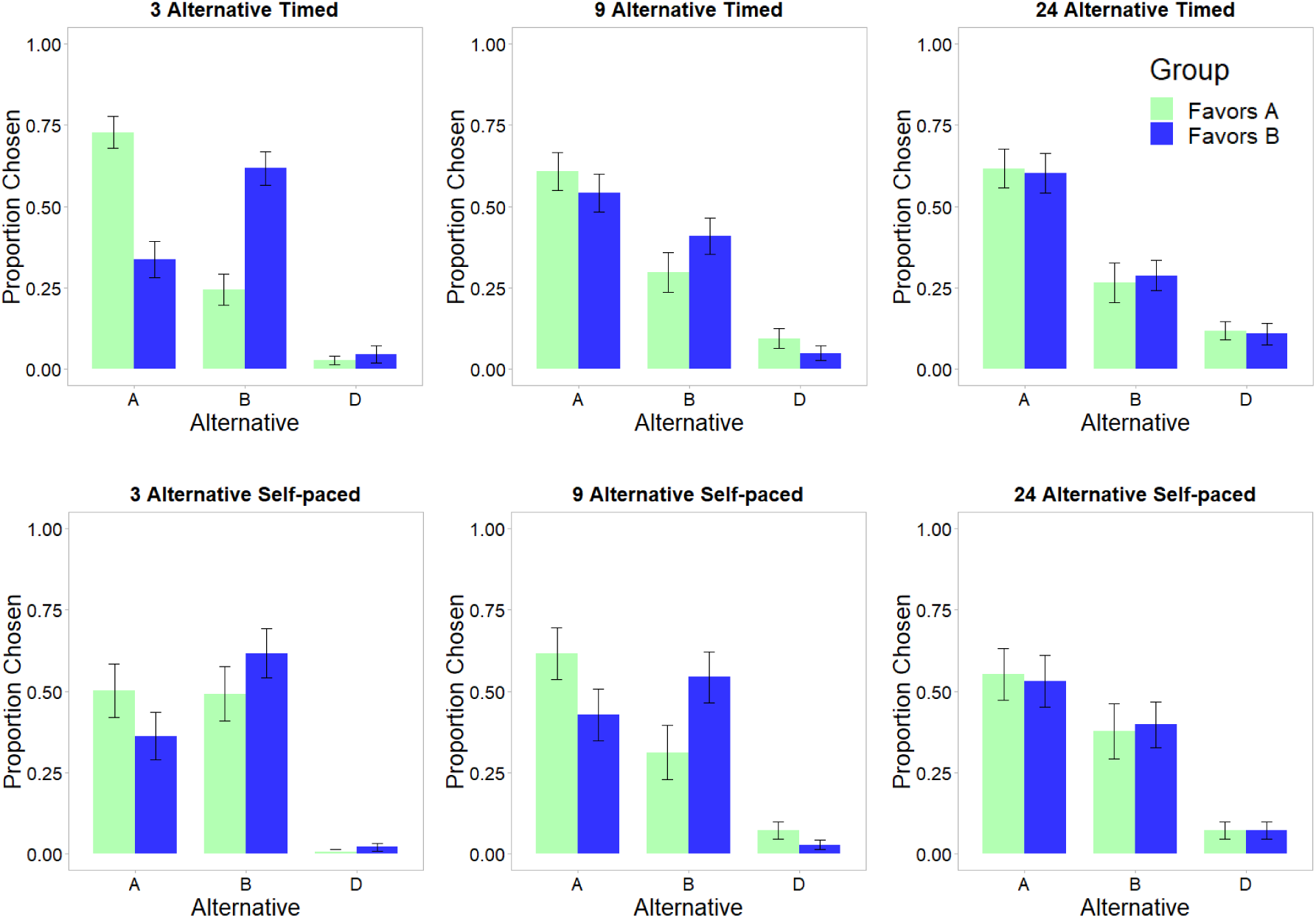
Choice Preference Shifts for Behavioral Experiment 2

Table 1 displays the average judgement ratings for each experimental condition for the three rating scales. We conducted a three-way Number x Decoy x Timing ANOVA on satisfaction ratings. A main effect of Number revealed that choices made from decisions with 24 options were less satisfying than those from three and nine options, *F*(1, 51) = 15.05, *p* < 0.001, η^2^_p_ = 0.23, reflected in the mean satisfaction of 6.59, 6.74, and 6.18 for three, nine, and 24 option choice sets, respectively.

**Table 1.**
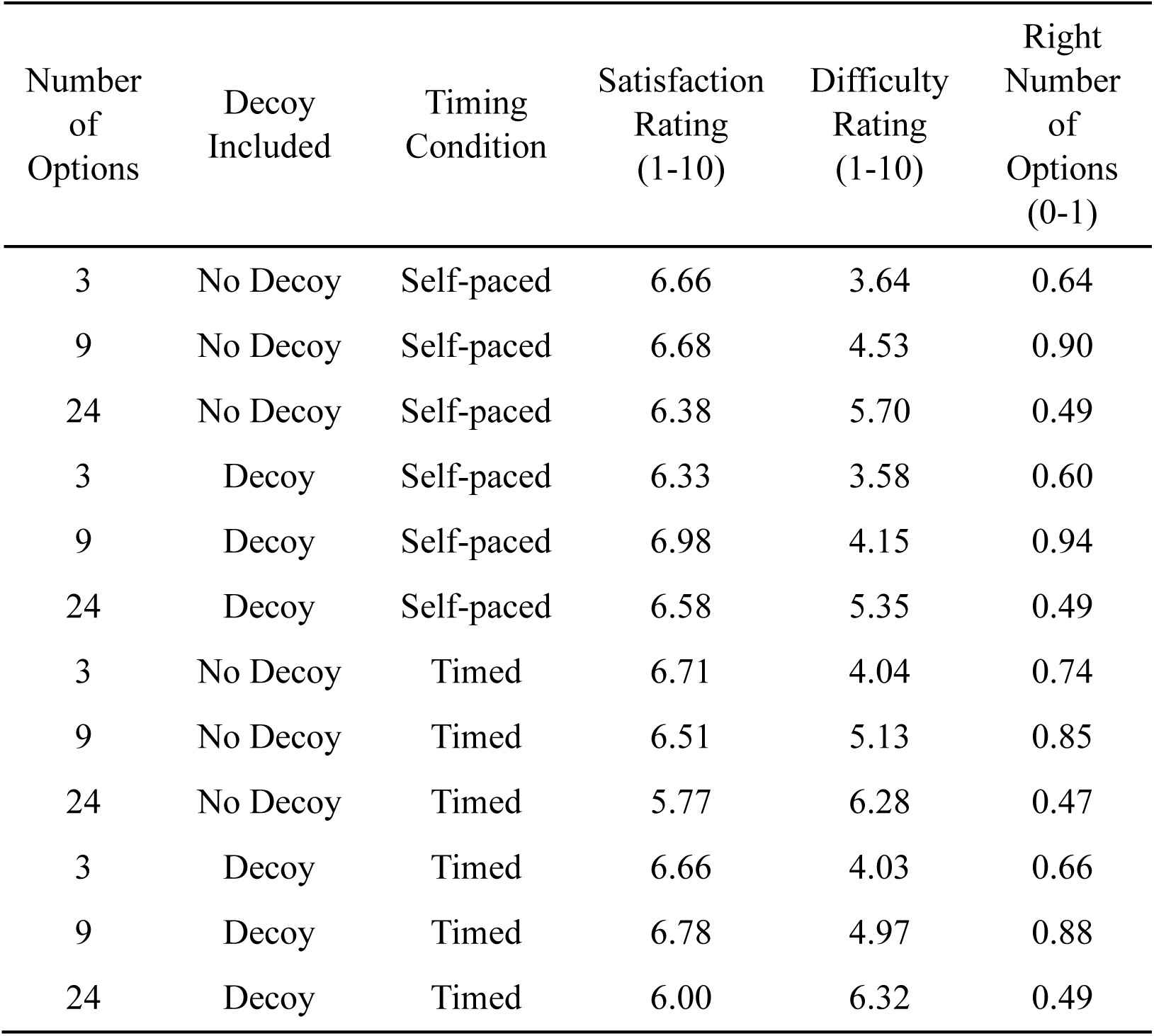
Mean Judgement Ratings for Behavioral Experiment 2.

| Number of Options | Decoy Included | Timing Condition | Satisfaction Rating (1-10) | Difficulty Rating (1-10) | Right Number of Options (0-1) |
| --- | --- | --- | --- | --- | --- |
| 3 | No Decoy | Self-paced | 6.66 | 3.64 | 0.64 |
| 9 | No Decoy | Self-paced | 6.68 | 4.53 | 0.90 |
| 24 | No Decoy | Self-paced | 6.38 | 5.70 | 0.49 |
| 3 | Decoy | Self-paced | 6.33 | 3.58 | 0.60 |
| 9 | Decoy | Self-paced | 6.98 | 4.15 | 0.94 |
| 24 | Decoy | Self-paced | 6.58 | 5.35 | 0.49 |
| 3 | No Decoy | Timed | 6.71 | 4.04 | 0.74 |
| 9 | No Decoy | Timed | 6.51 | 5.13 | 0.85 |
| 24 | No Decoy | Timed | 5.77 | 6.28 | 0.47 |
| 3 | Decoy | Timed | 6.66 | 4.03 | 0.66 |
| 9 | Decoy | Timed | 6.78 | 4.97 | 0.88 |
| 24 | Decoy | Timed | 6.00 | 6.32 | 0.49 |

There was a significant two-way interaction of the number of options and timing conditions on satisfaction ratings, *F*(1, 51) = 7.53, *p* < 0.01, η^2^_p_ = 0.13. Participants in the timed condition rated choice sets with 24 options significantly less satisfying than choice sets with three options, *t*(29) = 3.48, *p* < .01, *d* = 0.90, and nine options, *t*(29) = 5.48, *p* < .001, *d* = 1.41, with no significant difference between three and nine. Participants in the self-paced condition rated choice sets with 9 options significantly more satisfying than choice sets with three options, *t*(22) = 2.35, *p* < .05, *d* = 0.69, and 24 options, *t*(22) = 3.32, *p* < .01, *d* = 0.98, with no significant differences between three and 24 options.

There was a significant two-way interaction of the number of options and decoy conditions on satisfaction ratings, *F*(1, 51) = 5.68, *p* = 0.02, η^2^_p_ = 0.10. Choice sets with 9 options were significantly more satisfying with dominant options included *t*(52) = 3.14, *p* < 0.01, *d* = 0.43.

We conducted a three-way Number x Decoy x Timing ANOVA on difficulty ratings. A significant main effect of Number, *F*(1, 51) = 110.13, *p* < .001, η^2^_p_ = .67, indicated that difficulty ratings increased significantly with the increase in the number of options, reflected in the mean ratings of 3.85, 4.74, and 5.97 for three, nine, and 24 option choice sets, respectively. There was also a significant main effect of Decoy, *F*(1, 51) = 4.78, *p* < .05, η^2^_p_ =.09, reflected in the mean difficulty ratings of 4.92 and 4.78 for no decoy and decoy, respectively. Post-hoc comparisons of difficulty rating differences due to decoys at level of option number revealed significant differences only in nine option choice sets, *t*(52) = 3.14, *p* < .01, *d* = 0.43, but not for choice sets with three and 24 options. A significant main effect of Timing, *F*(1, 51) = 6.54, *p* < .05, η^2^_p_ = 0.11, indicated that choices made from the timed conditions were rated as more difficult than the self-paced condition.

We conducted a three-way Number x Decoy x Timing ANOVA on right number of options ratings that have been converted to the 0 to 1 scale as in Behavioral Experiment 1. The main effect of Number, *F*(1, 51) = 26.85, *p* < .001, η^2^_p_ = 0.34, indicated that choice sets with nine options were rated as more optimal than choice sets with three and 24 options, reflected in the mean ratings of .65, .83, and .48 for choice sets with three, nine, and 24 options, respectively. A significant interaction of Number and Decoy, *F*(1, 51) = 4.49, *p* < .05, η^2^_p_ = 0.08, indicated that the effect of Decoy was different for each level of number of options. Three option choice sets with decoys were rated as less optimal than choice sets without decoys, *t*(52) = 2.81, *p* < .01, *d* = 0.55. Nine option choices sets with decoys were rated as more optimal than choice sets without decoys, *t*(52) = 2.23, *p* < .05, *d* = 0.43. There was no significant difference in 24 option choice sets.

### Discussion

The results of these two behavioral experiments provide establish the validity of the design used in our neuroimaging study. We find strong evidence that nine options were perceived as the ideal amount for our grocery item stimuli. Ratings of the right number of options reached their peak in both behavioral experiments at nine items. This peak was consistent regardless of the presence or absence of decoy options, and, critically, regardless of timeframe. Nine options was rated as ideal for both the timed and self-paced condition in the second behavioral experiment. This is an important consideration as participants are restricted in their deliberation time due to the limitations imposed by the fMRI scanner. Additionally, three items were consistently rated as too few and 24 items were rated as too many. We also saw that choices made from choice sets with 24 options were rated as significantly less satisfying compared to choice sets with three and nine options. This provides evidence that we are inducing choice overload in our large choice sets as one of the main effects of choice overload is reduced choice satisfaction (Chernev et al., 2015). These studies also demonstrate the efficacy of our stimuli for studying the contextual effects of dominance, particularly in smaller choice sets.

## Neuroimaging Study

The present study investigates the neural correlates of choice overload in value-based multiattribute decision making. To this end, we test five primary hypotheses:

1. The anterior cingulate cortex (ACC), the dorsolateral prefrontal cortex (DLPFC), and the dorsal striatum (caudate nucleus and putamen) will show greater activity when participants choose from choice sets of moderate (nine) size than when choosing from choice sets of small (three) and large (24) sizes (Reutskaja, 2018; Woo et al., 2021).
2. Reductions in choice overload through the manipulation of choice complexity will be reflected in an overall increase in activity of the ACC, DLPFC, dorsal striatum and anterior insula (AIns). Choice sets with dominating alternatives will elicit greater activity in these brain areas than choice sets without dominating alternatives (Busemeyer et al., 2019; Hedgcock & Rao, 2009; Hu & Yu, 2014; Mohr et al., 2017; Reutskaja, 2018).
3. More activity will be observed in the AIns for participants who frequently choose a dominating option, and more activity will be observed in the ACC for participants who frequently *do not* choose a dominating option (Hu & Yu, 2014).
4. Preferences will be greater for dominating alternatives over competing alternatives within subjects and this effect will be attenuated with increasing numbers of alternatives. Weighting of a single attribute will increase and dominance weighting will decrease with larger assortments. There will be an increase in the tendency to use a lexicographic decision strategy as the number of alternatives increases. (Huber & Puto, 1983; Simonson, 1989; Simonson & Tversky, 1992; Stanley & Wedell, 2024; Trueblood et al., 2013; Wedell, 1991).
5. Greater activity in the DLPFC and posterior parietal cortex and less activity in the ACC will be observed for participants who tend to engage in a lexicographic decision strategy more often. Participants who do not engage in a lexicographic strategy as often will show greater activity in the AIns and ventromedial prefrontal cortex. (Khader et al., 2011; Van Duijvenvoorde et al., 2016; Venkatraman et al., 2009; Wichary & Smolen, 2016)

### Methods

#### Participants

A total of 40 participants (age range, 18 – 36 years; mean age, 22.3 years; 14 males) from the University of South Carolina campus were recruited. Data from two participants were excluded before analyses because of excessive head movement (>3 mm), and an additional three participants were excluded due to poor data quality causing misalignment during coregistration, leaving 35 individuals in the final sample. Participants were given $29.10 upon completion of the study. All participants gave written informed consent under a protocol approved by the Institutional Review Board of The University of South Carolina.

#### Procedure

The fMRI task design and timing were similar to that used by Reutskaja et al. (2018). Participants began by signing a consent form and completing practice trials to familiarize them with the task. A 3T MRI scanner was used to acquire brain images. The functional MRI scans were divided into 72 blocks, with one trial per block, divided equally across 3 scanning sessions of 14 minutes each. Participants were instructed to imagine that they are shopping for groceries for their family online and that they should attempt to find the most attractive price and quality rating combination. This dependent variable has been shown to produce strong manipulations of context (Stanley & Wedell, 2024; Wedell et al., 2022) and is designed to motivate participants to consider both attributes in their decisions. Figure 6 illustrates the timeline for each trial. Each trial consisted of a 10 second initial baseline epoch with a central fixation, a 10 second initial exposure to the choice set, a 10 second delay with central fixation, and a 5 second response phase displaying the choice set a second time. During the response phase, a cursor became visible which signaled for participants to use the MRI compatible joystick to move the cursor and select their preferred alternative by pressing a button on top of the joystick. The 10 second fixations separate the exposure and response phases to isolate decision related activity from potential sources of noise such as motor movements required for responding. The presentation order of the trials was randomized for each participant. At the end of the imaging session, a high-resolution anatomical scan was acquired. After the imaging session, a short survey was administered with 18 trials following the same design as the fMRI task. After each trial, participants were asked to provide satisfaction, difficulty, and ‘right number of options’ ratings as was performed in the behavioral pilot to assess susceptibility to choice overload for each participant.

**Figure 6.**
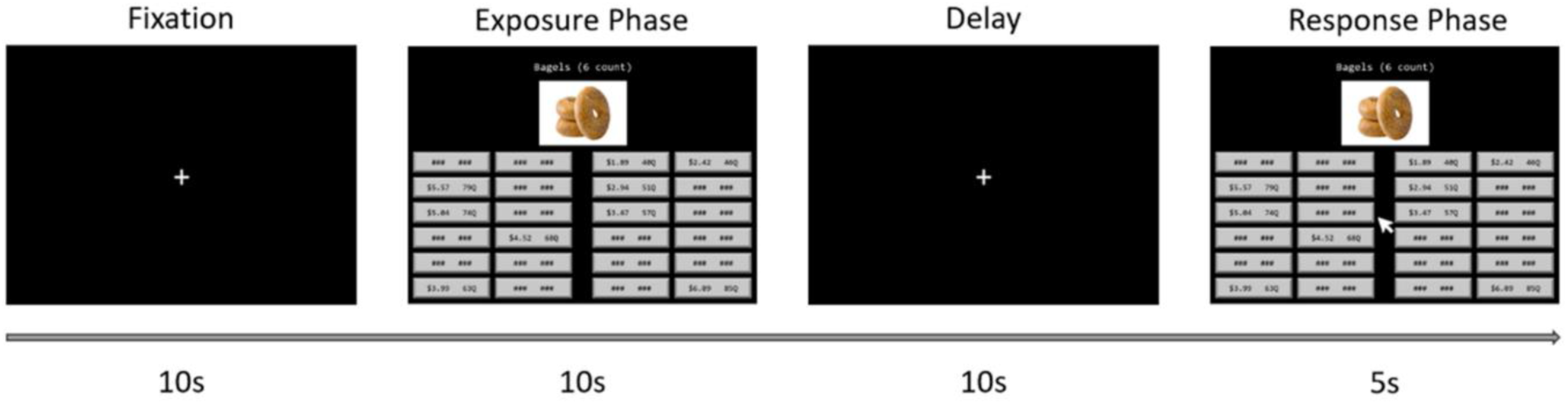
Trial Timeline for Neuroimaging Study

### Materials and Design

Stimuli were based on those used in Stanley and Wedell (2024) and Wedell et al. (2022), and they were presented using the E-Prime 3.0 software (Psychology Software Tools, Pittsburgh, PA). Thirty-six unique consumer items were presented to participants, repeated once each, for a total of 72 trials. For each presented stimulus, a number of options was available to choose from (either three, nine, or 24), each with a unique price ‘$’ and quality rating ‘Q’ along with a stock image and text description of the item (see Figure 7). All stimuli were presented with 24 buttons on the display in order to diminish visual differences between choice sets of different sizes (luminance, color, etc.).

**Figure 7.**
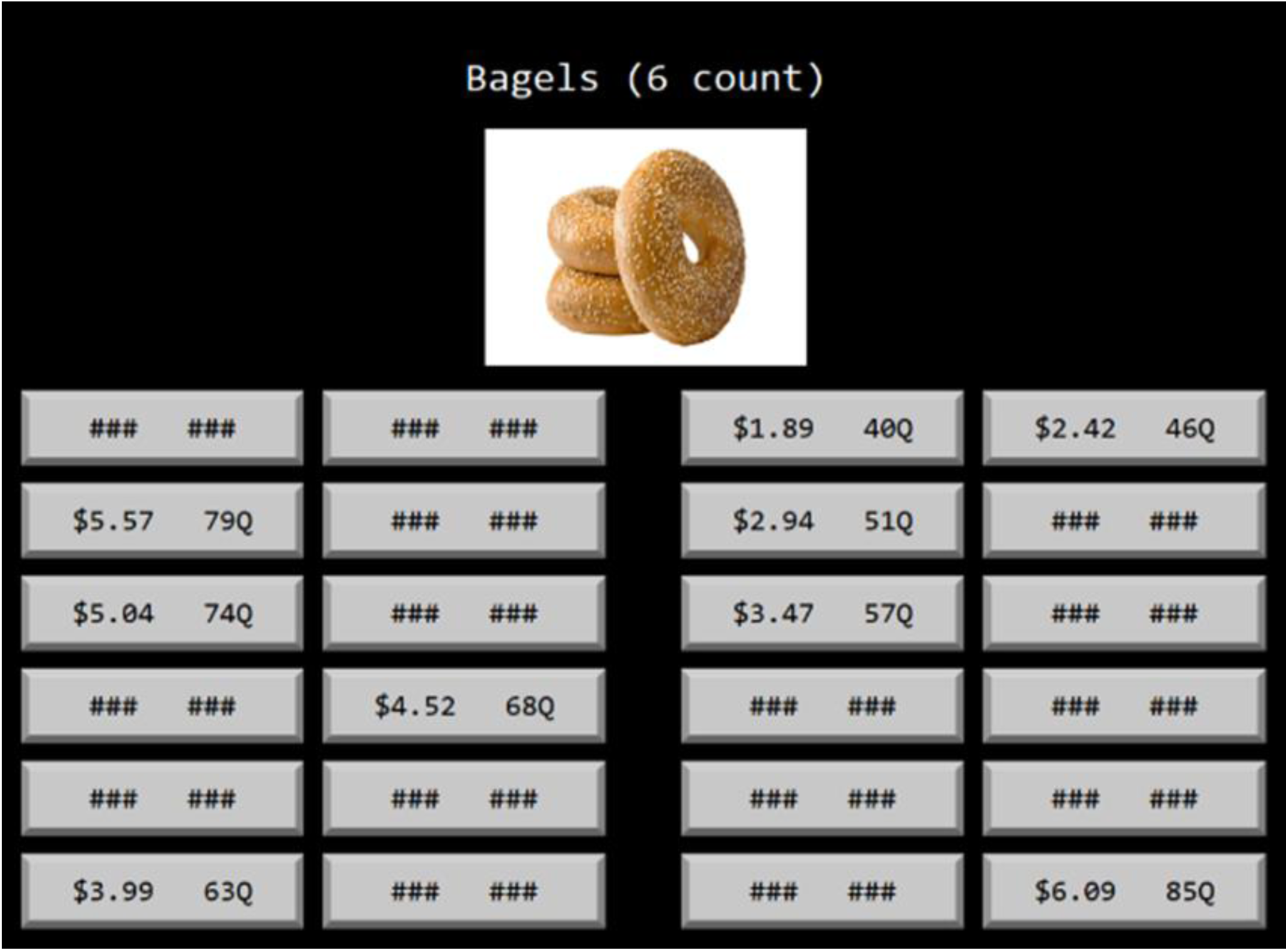
Example Stimulus Display

In choice sets with three and nine alternatives, the unused buttons were filled in with ‘###’ symbols, and participants were instructed not to select those options. The positions of the options were randomized within the 24 buttons, and the order of trials were randomized for each participant. Attribute values for alternatives were generated in the same manner as for Behavioral Experiment 2.

The stimuli were divided into six conditions: 3 (number of options: three, nine, 24) x 2 (dominating alternatives: included, not included). Assignment of the 72 consumer items to the six conditions were counterbalanced between subjects to mitigate potential biases from individual item preferences affecting decision strategies. The stimuli that include dominating alternatives were divided into three groups of alternatives of equal number on each trial as in

Stanley and Wedell (2024), group A, B, and D (see Figure 4). Group A and B traded off on price and quality competitively. Alternatives in group A were similar to one another and were relatively expensive and higher quality. Alternatives in group B were similar to one another and were relatively cheap and lower quality. Alternatives in group D were dominated by group A on half of the trials and dominated by group B on the other half (18 trials each). When dominated by group A, group D had similar quality ratings to group A but were more expensive. When dominated by group B, group D had similar prices to group B but had lower quality ratings.

Stimuli presented without including dominating alternatives were not divided into groups and had all options trade-off on price and quality competitively with one another such that prices of the options span the entire range of those surveyed at retailers. Figure 4 displays the relational structure of the alternatives within each condition.

#### Image Acquisition

Functional MRI data were acquired using a gradient-echo EPI sequence on a Siemens 3T scanner and 20-channel head coil at the McCausland Brain Imaging Center. Each participant performed three runs, each consisting of 568 volumes [repetition time (TR): 1.5 s; echo time (TE): 30 ms; voxel dimensions: 3 x 3 x 3 mm; 50 slices; 70 x 70 matrix; field of view (FOV): 210 mm; flip angle: 62°]. High-resolution T1-weighted anatomical images were acquired using a 3D inversion recovery-prepared gradient echo sequence (TR: 2300 ms; TE: 2.98 ms; voxel dimensions: 1 x 1 x 1 mm; 256 x 256 matrix; FOV: 256 mm; flip angle: 7°].

#### fMRI Data Processing and Analysis

We used SPM12 (Ashburner et al., 2014) to perform the functional image analyses. Image preprocessing was completed as follows. First, a mean image was created by realigning all functional images to the first scan of the first run. Next, the mean image of the realigned functional scans was co-registered to the anatomical image. The anatomical image was normalized to the SPM T1-template in Montreal Neurological Institute (MNI) space. The resulting nonlinear 3D transformation was applied to all EPI images. We then spatially smoothed the normalized functional images using a Gaussian kernel (6 x 6 x 6 mm3 full-width at half-maximum) and applied a high-pass filter (cutoff period 128ms). For our analysis we modeled the six experimental conditions separately (3 number of options (three, nine, 24) x 2 dominating alternatives (included, not included)) as well as six motion correction parameters obtained from a rigid-body transformation during image realignment.

### Results

#### Choice Data

To examine the effect of the decoys on choice preferences, Decoy Effect scores were calculated in the same manner as for Behavioral Experiment 2. These decoy effect scores were calculated for each participant and for each level of the number of options factor. There were large attraction effects for choice sets with three alternatives. However, there was no significant decoy effect for choice sets with nine and 24 options when averaging across the entire sample (Table 2).

**Table 2.** Decoy Effect by Number of Alternatives.

| Number of Alternatives | Choose Target | Choose Competitor | Choose Decoy | Decoy Effect |
| --- | --- | --- | --- | --- |
| 3 | 0.63 | 0.35 | 0.03 | 0.28*** |
| 9 | 0.51 | 0.44 | 0.05 | 0.06 |
| 24 | 0.44 | 0.45 | 0.11 | -0.01 |

\*\*\* $p \leq .001$

We conducted a three-way Number (3, 9, 24) x Favors (A, B) x Alternative (A, B) repeated measures ANOVA on choice proportions for A and B for trials that included decoy options. The Favors factor describes whether the decoy targets, or favors, the A or B alternative(s). The Alternative factor describes the two types of core alternatives, either high quality and high price (A) or low quality and low price (B). The Favors x Alternative interaction, *F*(1, 34) = 12.84, *p* < 0.001, 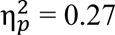, indicated significant context effects averaging across number of alternatives. The Favors x Alternative x Number interaction, *F*(2, 68) = 6.33, *p* < 0.01, 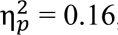, indicated a significant reduction in context effects as the number of alternatives increased. The main effect of Number, *F*(2, 68) = 19.11, *p* < 0.001, 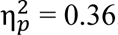, indicated that participants were more likely to choose the dominated decoy options as the number of alternatives increased. The Alternative x Number interaction, *F*(2, 68) = 4.62, *p* < 0.05, 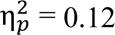, indicated that participants increased their preferences for high quality items over lower quality items as the number of options increased (Table 3).

**Table 3.** Attribute Preferences by Choice Set Size.

| Number of Alternatives | Group A Alternatives | Group B Alternatives |
| --- | --- | --- |
| 3 | .48 | .46 |
| 9 | .45 | .44 |
| 24 | .50 | .32 |

To test the hypothesis that the attraction effect is related to System 2 thinking, we computed Pearson correlations between participants’ scores on the Cognitive Reflection Test and their tendency to show the attraction effect. Contrary to our hypothesis, CRT scores were not significantly correlated with the attraction effect at any level of option numbers.

#### Post Scan Judgements

After the neuroimaging task, participants were asked to complete 18 additional trials that had them choose grocery items in a similar manner as they did while in the scanner. After each trial participants were asked three questions: “How satisfied are you with your choice?”, “How difficult was it to make your decision?”, and “Did you feel you had the right number of options to choose from?” Figure 8 displays the mean ratings from each of the six experimental conditions.

**Figure 8.**
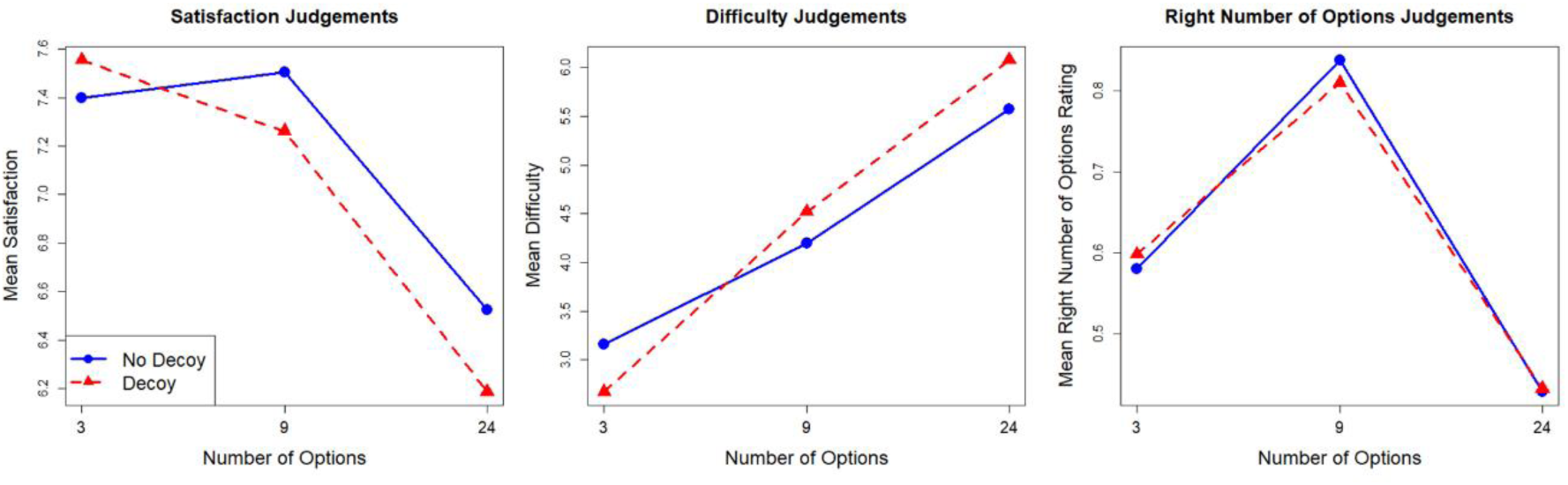
Judgement Ratings for Post-scan Survey

A two-way Number x Decoy repeated measures ANOVA revealed that satisfaction ratings differed significantly across choice sets with different numbers of alternatives *F*(2, 78) = 17.90, *p* < .001, 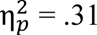. Participants were less satisfied with choices made from 24 options compared to choices made from three options, *t*(39) = 4.73, *p* < .001, *d* = 0.75, and compared to choices made from nine options, *t*(39) = 5.14, *p* < .001, *d* = 0.81. Satisfaction ratings from choices made from three and nine options were not significantly different. Additionally, there was no main effect of decoys on satisfaction ratings, nor any simple main effects of decoys at any level of option number.

A two-way Number x Decoy repeated measures ANOVA revealed that participants rated choices with more alternatives as more difficult, *F*(2, 78) = 61.68, *p* < .001, η_p_^2^ = .61. Difficulty ratings were significantly higher for choices with nine options compared to three options, *t*(39) = 7.013, *p* < .001, *d* = 1.11, and difficulty ratings were significantly higher for choices with 24 options compared to nine options, *t*(39) = 6.03, *p* < .001, *d* = 0.95. There was not a main effect of decoys on difficulty ratings; however, three option choice set with decoys were rated as less difficult than those without decoys, *t*(39) = 2.70, *p* = 0.01, *d* = 0.43, and 24 option choice sets with decoys were rated as more difficult than those without decoys, *t*(39) = 2.24, *p* < .05, *d* = 0.35. There was no simple main effect of decoy in choice sets with 9 options.

Ratings for the question “Did you feel you had the right number of options to choose from?” were given on a 1-9 scale where a 1 labeled as ‘Too few options’, 9 labeled as ‘Too many options’, and 5 labeled as ‘Just the right number of options.’ For interpretability, ratings were converted from this 1-9 scale a described in Behavioral Experiments 1 and 2. A two-way within-subjects ANOVA revealed that ratings for the right number of options differed significantly across the number of alternatives, *F*(2, 78) = 45.09, *p* < .001, η_p_^2^ = .54. Choice sets with 9 options were perceived as being more optimal than choice sets with three options, *t*(39) = 5.69, *p* < .001, *d* = 0.90, and more optimal than choice sets with 24 options *t*(39) = 10.59, *p* < .001, *d* = 1.67. Choice sets with three options were perceived as more optimal than choice sets with 24 options, *t*(39) = 3.42, *p* < .01, *d* = 0.54. There was no main effect of decoys, nor were there any simple main effects at any number of options level.

#### Computational Modeling

We modeled the choice data from the task performed in the scanner using a value-added approach (Pettibone & Wedell, 2000; Wedell, 1991; Wedell & Pettibone, 1996). We included parameters capturing a basic weighted additive model augmented with contextually sensitive parameters capturing dominance valuing and distance valuing. The general value function was as follows:

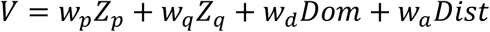

where 𝑍_𝑝_ is the z-score of price (across all price values in the experiment) weighted by 𝑤_𝑝_ and 𝑍_𝑞_ is the z-score of quality (across all quality values in the experiment) weighted by 𝑤_𝑞_. These first two terms of the equation represent a weighted additive model that is contextually insensitive. Note that we reverse the sign of 𝑍_𝑝_ so that a higher weighting for price reflects a preference for lower priced items.

The attraction effect is based on manipulating dominance asymmetrically. To capture dominance valuing, we assign 𝐷𝑜𝑚 a value of 1 if it dominates another alternative and a value of −1 for if an alternative dominates it, with 𝑤_𝑑_ representing the weight of this process. 𝐷𝑖𝑠𝑡, calculated as the city-block distance from the average of values in the contextual set for that trial. When 𝐷𝑖𝑠𝑡 is negatively weighted by 𝑤_𝑎_, there is an advantage of being close to the contextual average, enhancing the attraction effect (Wedell & Pettibone, 1999).

We used a proportional response function:

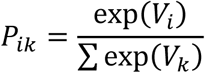

so that the probability of choosing alternative *i* in context *k* is proportional to the exponential value of *i* divided by the sum of the exponential values of the alternatives in the choice set. Model parameters shown in Table 4 are averages of the parameters fit for each participant for each level of option number.

**Table 4.** Computational Modeling Results.

| Number of Alternatives | CV LL | BIC | $W_P$ | $W_Q$ | $W_{Dom}$ | $W_{Dist}$ |
| --- | --- | --- | --- | --- | --- | --- |
| 3 | 654 | 2173 | 0.83 | 0.58 | 0.87 | -0.38 |
| 9 | 2142 | 6209 | 0.65 | 0.54 | 0.61 | -0.44 |
| 24 | 4400 | 11913 | 0.51 | 0.56 | 0.42 | -0.28 |

*Note.* Values in the CV LL column show the loglikelihood measures from the twofold cross validation procedure. A more negative $W_{Dist}$ indicates a greater preference for alternatives close to the contextual average.

Model parameters were fit for each number of alternatives using a Bayesian maximum a posteriori estimation method with a normal prior distribution possessing a mean of 0 and a standard deviation of 1. Model performances were assessed using 2-fold cross validation as well as by assessing the Bayesian information criteria (BIC) for each. The models were trained individually for each subject over a maximum of 500 iterations using the ‘optim’ function in R with the Nelder-Mead optimizer.

#### DLPFC shows Inverse U-shape activation as a function of choice set size

To investigate the relationship between brain activity within our regions of interest and choice set size, we tested two orthogonal hypotheses. Our primary hypothesis, in accordance with results from Reutskaja et al. (2018), predicted a quadratic relationship between choice set size and BOLD signal in the ACC, DLPFC, and dorsal striatum. Secondarily, we tested for linear relationships between choice set size and the same ROIs. Linear and quadratic contrasts were tested in separate general linear models (GLMs), averaging across the levels of the second experimental factor (decoy vs no decoy). Second-level analyses were restricted to voxels within anatomically defined a priori ROIs using explicit masks from the Neuromorphometrics atlas (Neuromorphometrics, Inc., 2012). Small volumes correction was applied, and significance was assessed at the voxel level using family-wise error correction within each ROI.

We identified two significant clusters within the DLPFC for the quadratic contrast, indicating greater activity for choice sets with 9 alternatives compared to choice sets with three and 24 alternatives (Table 5). For illustration, Figure 9 displays significant voxels from our ROI analysis that were significant at *p* < .001, uncorrected.

**Figure 9.**
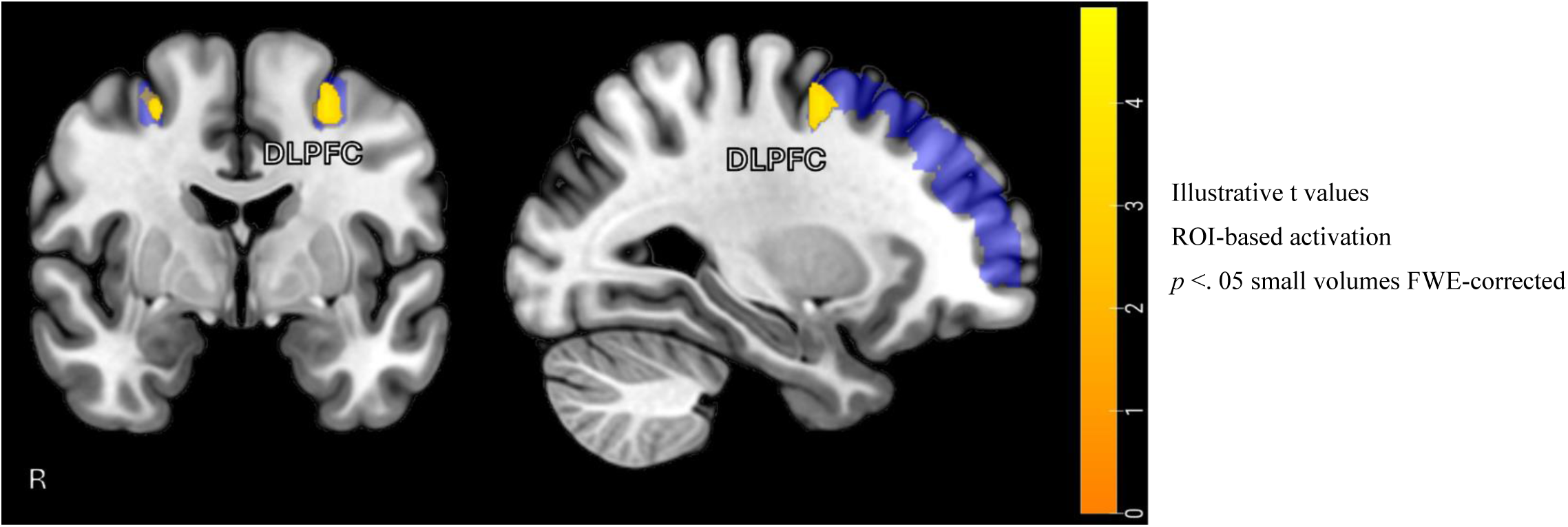
ROI-based Results for Quadratic Contrast. Areas highlighted in blue represent the Neuromorphometric anatomical atlas used for statistical analysis. Areas highlighted in orange show the locations of significant clusters. Illustrative t values are used.

**Table 5.** Significant Clusters for Quadratic Contrast.

| Structure | x, y, z<br>(mm MNI) | t-value<br>(peak value) | p<br>(FWE-corr.) | Cluster Size<br>(voxels) |
| --- | --- | --- | --- | --- |
| DLPFC | 27, -1, 56 | 5.11 | .013 | 14 |
| DLPFC | -24, -1, 53 | 4.86 | .025 | 29 |

For the linear contrast, we identified two significant clusters showing a positive linear relationship between DLPFC activity with choice set size (Table 6). We also identified a marginally significant cluster within the ACC. For illustration, Figure 10 displays significant voxels from our ROI analysis that were significant at *p* < .001, uncorrected.

**Figure 10.**
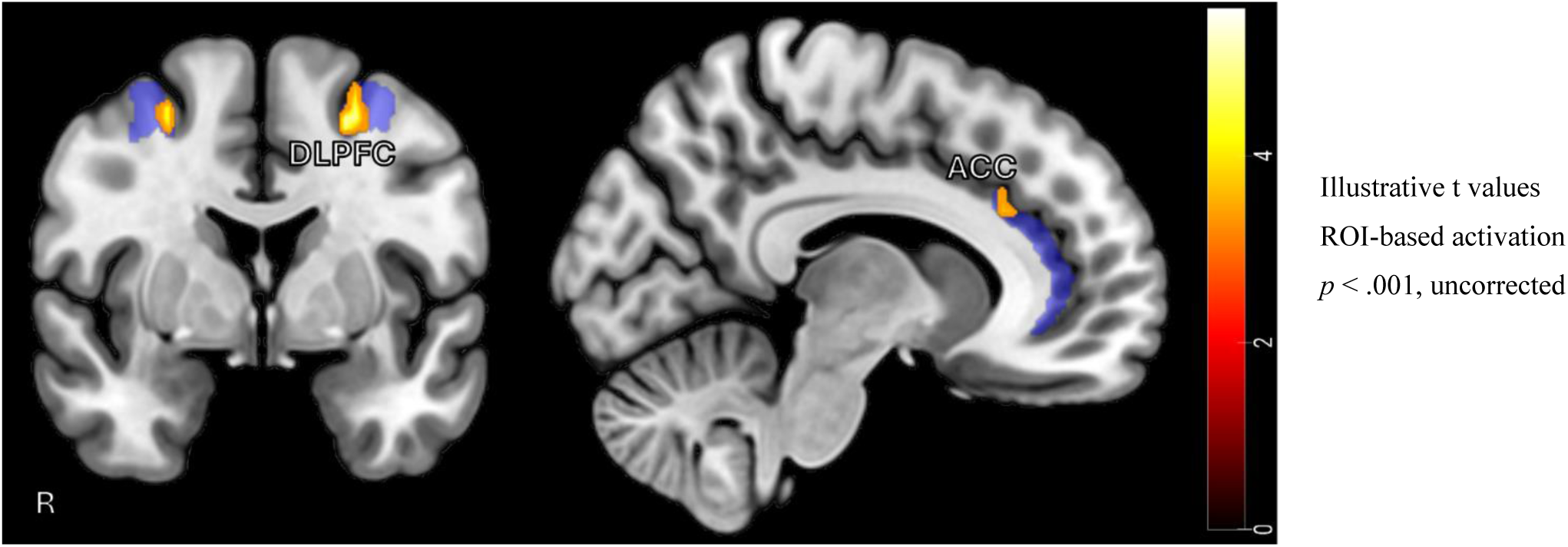
ROI-based Results for Linear Contrast. Areas highlighted in blue represent the Neuromorphometric anatomical atlas used for statistical analysis. Areas highlighted in orange show the locations of significant clusters. Illustrative t values are used.

**Table 6.** Significant Clusters for Linear Contrast.

| Structure | x, y, z<br>(mm MNI) | t-value<br>(peak value) | p<br>(FWE-corr.) | Cluster Size<br>(voxels) |
| --- | --- | --- | --- | --- |
| DLPFC | -27, 2, 56 | 5.65 | 0.003 | 32 |
| DLPFC | 27, -1, 53 | 4.93 | 0.023 | 12 |
| ACC | 9, 29, 26 | 3.99 | .051 | 9 |

*Note.* All statistical inference is based on family-wise error correction within anatomically defined ROIs using small volume correction in SPM12. ROIs were defined using the Neuromorphometric atlas distributed with SPM12. Coordinates are reported in MNI space.

#### The presence of decoy options increases DLPFC and AIns activity

To investigate the effect of choice set complexity on brain activity in our regions of interest, we examined whether BOLD activity differed between trials with decoy options and those without, collapsing across set size. First-level contrast images reflecting the comparison of decoy-present > decoy-absent trials were entered into a second-level one-sample t-test. Statistical inference was restricted to a priori regions of interest, including the ACC, DLPFC, putamen, caudate, and anterior insula, defined anatomically using the Neuromorphometrics atlas distributed with SPM12. Small volumes correction was applied, and significance was assessed at the voxel level using family-wise error correction within each ROI. We identified clusters within the DLPFC and the AIns that showed significantly greater activity to trials with decoys compared to trials without decoys (Table 7). For illustration, Figure 11 displays significant voxels from our ROI analysis that were significant at *p* < .001, uncorrected.

**Figure 11.**
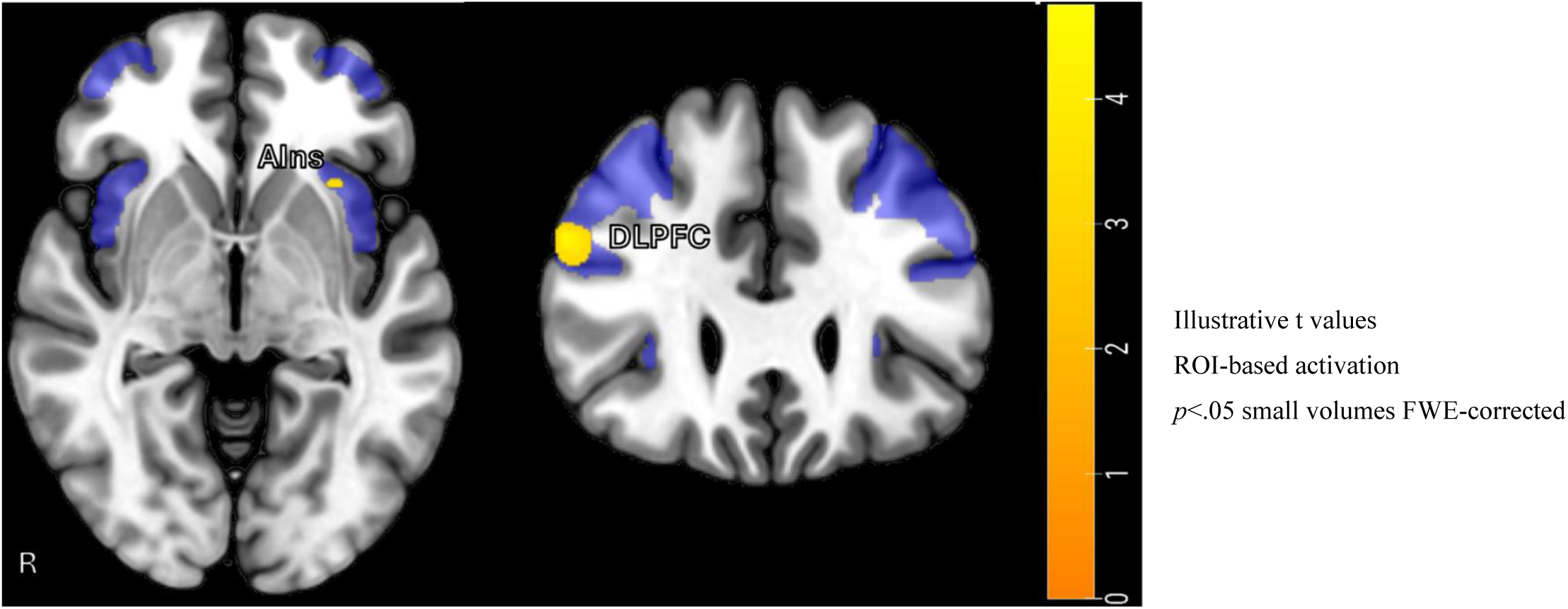
ROI-based Results for Decoy > No Decoy Contrast. Areas highlighted in blue represent the Neuromorphometric anatomical atlas used for statistical analysis. Areas highlighted in orange show the locations of significant clusters.

**Table 7.** Significant Clusters for Decoy >.

| Structure | x, y, z<br>(mm MNI) | t-value<br>(peak value) | p<br>(FWE-corr.) | Cluster Size<br>(voxels) |
| --- | --- | --- | --- | --- |
| DLPFC | 54, 29, 29 | 4.91 | 0.023 | 36 |
| AIns | 30, 14, -16 | 4.07 | 0.042 | 5 |

#### Activation in the ACC reflects dominance valuing

We analyzed how parameters derived from computationally modeled choice data relate to brain activity within our regions of interest. We fit our model to the choice data and extracted beta weights from first-level contrast images reflecting the comparison of decoy-present > decoy-absent trials for choice sets with 9 and 24 alternatives. We found a significant negative correlation, *r*(33) = -0.37, *p* < 0.05, between our parameter measuring dominance valuing and the brain activity measured in the ACC. In 9 and 24 option choice sets, participants with higher ACC engagement valued dominance less. This relationship is displayed in Figure 12.

**Figure 12.**
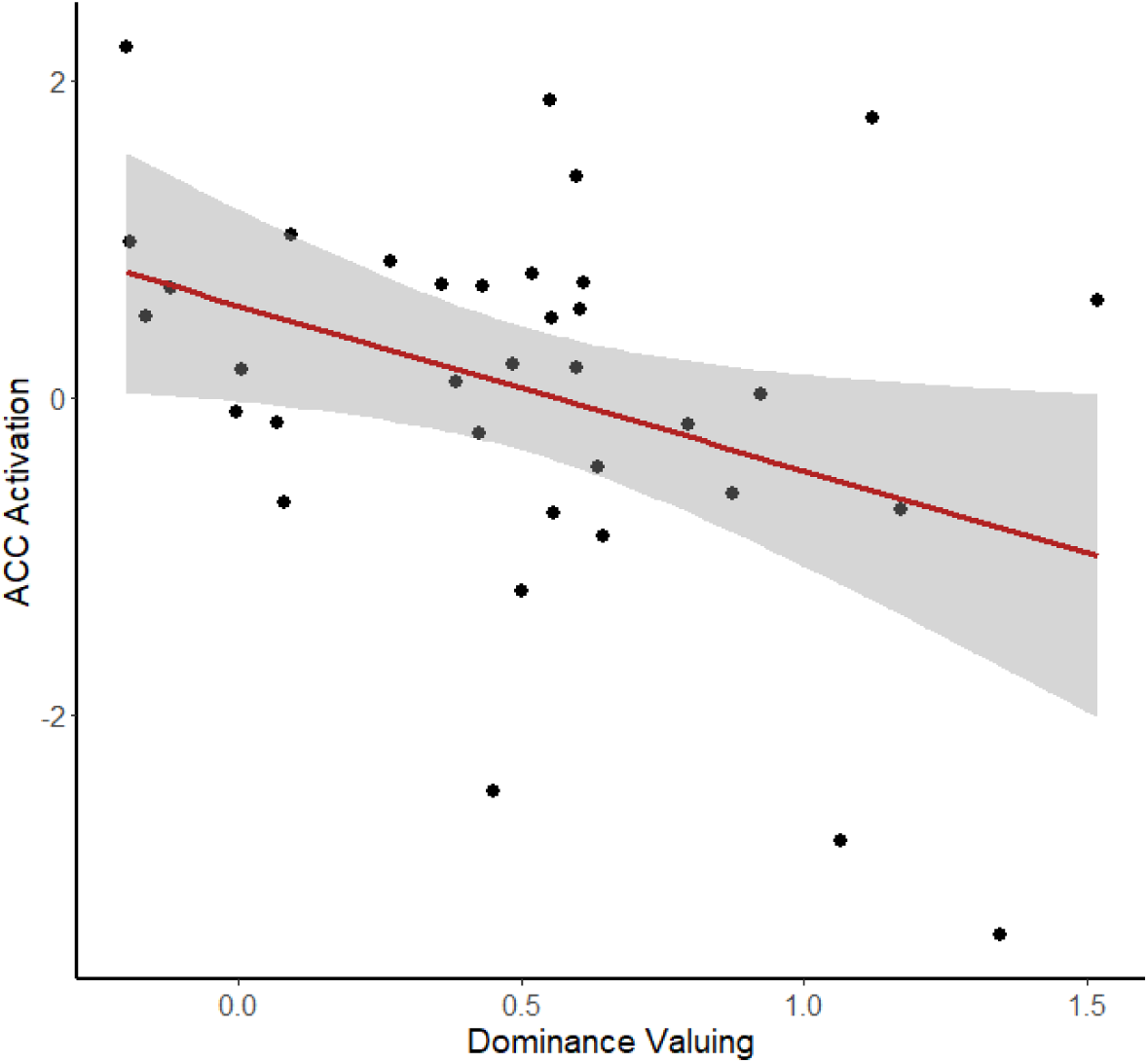
Correlation of Mean ACC Activity and Dominance Valuing

#### Activation in the ACC and AIns reflects decision strategy

In order to examine how brain activity relates to decision strategy, we separated our sample into two groups based on their tendency to utilize a lexicographic decision strategy. Participants were assigned to the Lex group if they chose the same type of item (either expensive and high quality or cheap and low quality) in more than 85% of choices with either 9 or 24 options. Otherwise, participants were assigned to the Non-Lex group. Beta weights were extracted from first-level contrast images reflecting the comparison of decoy-present > decoy-absent trials. Beta weights were significantly higher for the Non-Lex group compared to the Lex group in the ACC, *t*(33) = 2.58, *p* < 0.05, *d* = 0.90 and the AIns, *t*(33) = 2.09, *p* < 0.05, *d* = 0.73. Participants who tended *not* to utilize a simplifying lexicographic decision strategy showed greater difference in activity in the ACC and AIns in trials with decoys compared to trials without decoys.

#### Quadratic activity in the ACC and DLPFC reflects Grit-S effort scores

To examine if self-reported grit levels are reflected in brain activity, we correlated scores from the Grit-S with beta weights extracted from first-level contrast images reflecting a quadratic relationship with choice set size. We found a significant positive correlation between the effort construct of the Grit-S survey and beta weights extracted from both the ACC, *r*(33) = .34, *p* < .05, and the DLPFC, *r*(33) = .35, *p* < .05. Participants with a larger difference in BOLD signal between nine options and three and 24 options in the ACC and DLPFC reported greater levels of effort on the Grit-S survey. These relationships are displayed in Figure 13.

**Figure 13.**
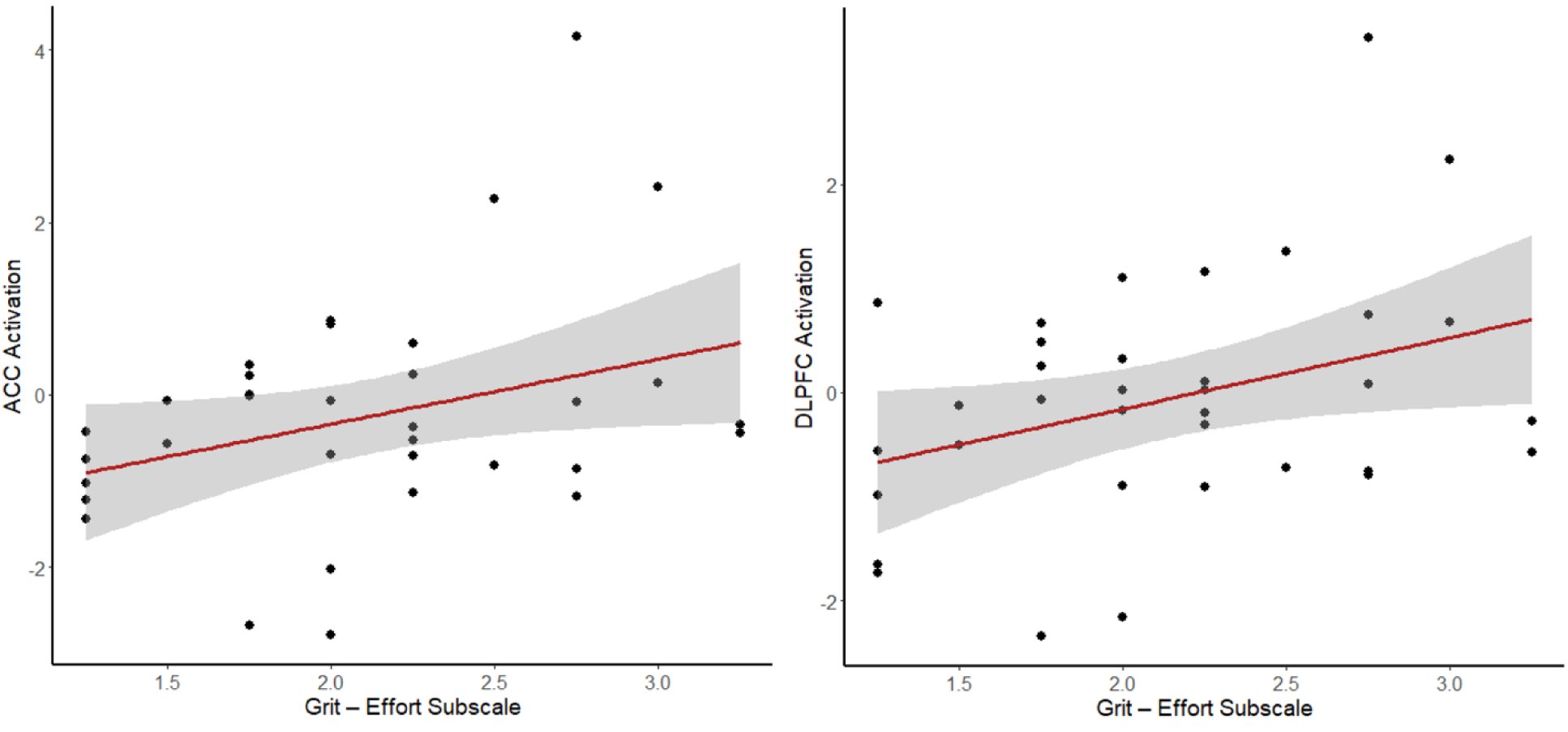
Correlations of Quadratic Brain Activity with Grit-S Effort Scores

## Discussion

### Hypothesis 1

The present study provides supporting evidence for activity in the ACC, DLPFC, and AIns as markers for choice overload. Choice overload is a mental state resulting from decision demands that exceed the available cognitive resources. Coombs and Avrunin’s (1977) theory of preference explains how we tend to perceive choice sets of moderate size as the most valuable, thus we should expect participants to be the most motivated to engage in cognitive control for choice sets with just the right number of options compared to choice sets that are too small (less need for cognitive control) and too large (choice overload resulting from excessive cognitive demands). As choice overload is characterized by decreased engagement with the choice process, we hypothesized brain areas involved in cognitive control should show greater activity in choice sets of moderate size (9) compared to choice sets that are too small (3) and too large (24). Our neuroimaging analysis found significant clusters within the left and right DLPFC that respond more strongly to choices with nine alternatives compared to choice sets with three and 24 alternatives. These results are in accordance with previous research from Reutskaja et al. (2018) that also identified a quadratic activity profile as a function of choice set size within the DLPFC.

The DLPFC has been shown to be involved in filtering irrelevant information. Woo et al. (2022) demonstrated that enhancement of DLPFC function using HD-tDCS attenuates the effect of attention duration to poor options on suboptimal choice behavior.

These results also agree with an interpretation of DLPFC function from MacDonald et al. (2000) that disassociates DLPFC and ACC function within cognitive control. In their study, they find that the DLFPC is responsible for the implementation of control, that is, maintaining the attentional demands of the task, whereas the ACC is responsible for conflict and performance monitoring. Our results provide converging evidence for the role of the DLPFC in filtering irrelevant information and the implementation of cognitive control during value-based decision making. The quadratic function of activity supports the interpretation that the DLPFC is maximally engaged when cognitive resources must be allocated efficiently, such as when the number of available options is optimally valuable and requires suppression of distractions and sustained goal maintenance. This pattern is consistent with the DLPFC’s role in top-down cognitive control processes for task relevant information.

We secondarily tested our regions of interest for significant linear relationships with choice set size. We found significant clusters within the DLPFC as well as a marginally significant (*p* = .051 small volumes FWE-corrected) within the ACC. A likely explanation for why the ACC is showing a trend more resemblant of a linear relationship rather than the quadratic relationship identified by Reutskaja et al. (2018) is the difference in stimuli and task design. Choosing a favorite landscape, as in Reutskaja et al. (2018), is a perceptual based task that likely engages holistic, intuitive valuation processes in addition to cognitive control. Participants were likely able to rely on predefined personal preferences (e.g., preference for beaches over forests) to guide their choices, especially because the participants in their study had previously seen the landscape stimuli in their pre-scan rating task. In contrast, the present study utilizes numeric choices with explicitly defined quantitative attributes that likely engage a more deliberate, utilitarian, or rules-based approach to the decision-making process. Additionally, Reutskaja et al. (2018) identified dorsal striatum activity that showed similar effects to the ACC, reflecting the reward potential of the choice sets. We were not able to replicate this activity within the dorsal striatum, so it could be that the design of the present study produced a weaker emotional or hedonic reward component, and that ACC activity in our study more so reflects effort, which grew with set size, rather than reward signal.

This ACC function distinction from the DLPFC provided by MacDonald et al. (2000) is mostly supported by the results of the present study. We find greater ACC activity for participants with less dominance valuing, suggesting the implementation of cognitive control through the ACC is required to deal with the conflict generated from resisting a heuristic style of decision making through choosing the dominating alternative. However, we would then expect to see significant clusters within the ACC for the decoy > no decoy contrast, which we did not. A possible explanation for this discrepancy could be high levels of individual differences. We may be testing across two or more subpopulations simultaneously, for example, those who attend to dominance and choose the target (middle – high ACC usage), those who attend to dominance and resist choosing the target (high ACC usage), and those who do not attend to dominance (low ACC usage). The dominance valuing parameter derived from our computational modeling is able to capture differences between these populations with more nuance than the decoy effect because the parameter weighs targets as 1, competitors as 0, and decoys as −1. In this way, the dominance valuing parameter is penalized in individuals who choose decoys, whereas the decoy effect only accounts for the relative choice proportions of targets and competitors. We find further evidence for this explanation in our analysis of how decision strategies are reflected in brain activity. Participants who tended not to engage in simplifying lexicographic strategies showed greater differences in the ACC for the decoy > no decoy contrast. Thus, for those participants that are using a compensatory approach and are more likely to notice dominance relationships, the ACC appears to be more engaged, reflecting the increased conflict monitoring demands associated with resisting heuristic-based choices. This supports the notion that the ACC becomes particularly active when participants must navigate competing response tendencies. Taken together, these findings refine the functional dissociation of the DLPFC and ACC. The DLPFC supports active maintenance and filtering required for efficient decision processing, while the ACC is sensitive to decisional conflict and effort demands of the choice environment.

We see a significant linear trend in the DLPFC in addition to the quadratic trend. This is likely due to the contrast designs that we use for our two hypotheses. The quadratic contrast averages activity from three and 24 alternative choice sets and compares them to choice sets with nine alternatives. We define the quadratic contrast as [−1 2 −1 −1 2 −1], collapsing across the decoy conditions. The linear contrast effectively shows brain activity that is greater in trials with 24 alternatives compared to trials with three alternatives. We define the linear contrast as [−1 0 1 - 1 0 1]. This allows for both a quadratic and a linear relationship within the DLPFC because, while DLPFC activity is highest in trials with nine options, it is more active in choice sets with 24 options compared to choice sets with three options. This finding is also consistent with the pattern identified in Reutskaja et al., (2018).

### Hypothesis 2

The present study finds supporting evidence for greater engagement of the DLPFC and AIns for decisions in choice sets that have decoy options compared to choice sets where all options trade-off equally on their attributes. These results are in agreement with previous research that finds similar effects. Hedgcock and Rao (2009) found greater DLPFC activity for choices with decoys compared to without decoys. Mohr et al. (2017) and Hu and Yu (2014) both found greater activity in the anterior insula for choice sets with decoys compared to without decoys. Reutskaja et al. (2018) explains the increase in DLPFC for choice sets with dominating alternatives as a marker of an increased value of the choice set size. Although a heuristic response to the decoy allows for a reduction in processing costs, the decoy increases the perceived attractiveness of the choice set and increases motivation to engage in cognitive control to maximize the expected value of the choice outcome. The AIns likely increases is activation to facilitate a heuristic, “gut feeling” response to the decoy. This result provides evidence for the role of the AIns in driving an affect component of decision making in response to the salience of the dominance relationship, possibly driven by an aversion to the strictly less valuable decoy option. This evidence is strengthened by our finding that participants who tended not to engage in a lexicographic decision strategy showed a greater difference in AIns and ACC activity compared to those who did. A lexicographic decision strategy that attempts to maximize on a single attribute fails to consider many of the available options in a choice set, reducing the likelihood that they recognize a dominance relationship within the choice set.

### Hypotheses 3 and 4

We did not find direct support for our third hypothesis that predicted the linear relationships between decoy effect and brain activity in the ACC and AIns. However, we did find correlations between our dominance valuing parameter derived from the computational modeling of choice behavior. As mentioned earlier, we explain this discrepancy through high individual differences. While there was a strong attraction effect in choice sets with three alternatives, choice sets with nine and 24 alternatives generated strong repulsion effects in some participants in our sample. In total, at the group level, we observed no decoy effect on average in choice sets with nine and 24 alternatives. We also found that participants were more likely to engage in a lexicographic decision strategy as the number of options increased, which reduces the probability that dominance relationships are noticed. This was reflected in an increase in preference for expensive, high quality items as the number of options increased.

### Hypothesis 5

Hypothesis 5 looked at how different decision strategies, inferred from choice behavior, related to brain activity. While we did not find significant differences in the DLPFC, posterior parietal cortex, or the VMPFC, we did see an effect within the ACC and AIns. Participants who avoided using a lexicographic decision strategy showed greater mean beta weights extracted from the Neuromorphometric atlas anatomical ROIs for the decoy > no decoy contrast compared to participants who did use the simplifying strategy. We interpret these results as further linking the ACC to cognitive control and effort, and the increase in AIns activation is likely a result of a compensatory strategy causing a greater likelihood of noticing the dominance relationship between targets and decoys. These results show how decision strategies are differentially represented in the brain.

### Exploration of Individual Differences Measures

Participants completed the cognitive reflection test (CRT) and the Grit-S questionnaire during the post-scan survey so that we could investigate some exploratory hypotheses relating to these individual difference measures. We predicted that scores on the CRT would be correlated with the decoy effect, as we hypothesized that the attraction effect is driven by System 1 thinking (Kahneman, 2011). However, we failed to find any evidence for this relationship. We also tested correlations between the effort construct of the Grit-S questionnaire and ACC activity. We found significant correlations between the effort construct of the Grit-S survey and mean beta weights from the quadratic contrast within the ACC and DLPFC. These results provide supporting evidence for their role in cognitive control. We find that participants who self-reported as putting in more effort into their daily activities showed greater differences in ACC and DLPFC activity for trials with nine alternatives compared to trials with three and 24 alternatives. These results reflect how individual differences shape how the brain differentially responds to the context of the choice set and the amount of effort one puts into the decision process. These results also support previous research that shows the ACC is a central hub for the brain representation of grit and tenacity (Touroutoglou et al., 2020). A description of an integrated neural model of value-based, multiattribute decision making is presented in the Supplement.

### Conclusion

Together, these findings provide evidence that choice overload in multiattribute decision making reflects coordinated, but dissociable, contributions of cognitive control, conflict monitoring, and salience systems. The DLPFC exhibited a single-peaked engagement profile aligned with optimal assortment size, while the ACC and AIns tracked the escalating effort and conflict associated with large and complex choice environments. By integrating fMRI analyses with computational modeling and behavioral judgements across varying assortment sizes, this study demonstrates how decision strategies shift as demands increase and how these shifts are expressed in target neural circuits. These results advance a mechanistic account of how the brain evaluates and navigates complex value-based decision environments, bridging literature on context effects, cognitive control, and consumer choice. More broadly, the results highlight that modern decision environments, characterized by abundant options, place meaningful constraints on human cognition suggesting avenues for designing choice architectures that better align with the limits of the neural systems that support everyday decision making.

