## Supplemental Materials for "Neural Dynamics of Multiattribute Decision Making Under Choice Overload"

**Stimuli Creation**

For each grocery item that was used, price and quality values were created as follows. A researcher visited multiple grocery stores and recorded the highest priced and lowest priced option available for a given grocery item. From these two values, a price average and a price range were created for each item.

For choice sets with a single equi-preference contour, prices were calculated for items as follows. For each grocery item and for each number of options for choice set, a price step size was created such that:

$$P_{Step} = \frac{P_{High} - P_{Avg}}{\left\lfloor \frac{N}{2} \right\rfloor}$$

where $P_{Step}$ is the price difference between adjacent items along the equi-preference contour, $P_{Avg}$ is the average price of the item, $P_{High}$ is the upper range value of the item, and $N$ is the number of options in the choice set. The floor function ensures that when $N$ is odd, price steps are distributed symmetrically around the average by assigning equal spacing to items on either side of the midpoint. Prices were then generated for each option, in decreasing order, such that:

$$P_i = P_{Avg} + (P_{Step} * \left( \left\lceil \frac{N}{2} \right\rceil - i \right))$$

where $P_i$ is the price of item $i$, and $i$ ranges from 1 (the most expensive item) to $N$ (the least expensive item), and where $N$ is the number of options in the choice set. This makes it so that the price of each item is equally spaced from one another and that the most extreme items have price values equivalent to the upper and lower range values of that particular grocery item.

Quality values for each grocery item were generated to fall between 40 and 100. Low quality values and high quality values were created using random integers ranging between 40 to

50 and 90 to 100, respectively. These high and low values were used to compute an average quality and a quality range for each grocery item. Similar to prices, for each grocery item and for each number of options, a quality step was created such that:

$$Q_{Step} = \frac{Q_{High} - Q_{Avg}}{\left\lfloor \frac{N}{2} \right\rfloor}$$

where $Q_{Step}$ is the quality difference between adjacent items along the equi-preference contour, $Q_{Avg}$ is the average quality of the item, $Q_{High}$ is the upper range value of the item, and $N$ is the number of options in the choice set. This makes it so that the quality of each item is equally spaced from one another and that the most extreme items have quality values equivalent to the upper and lower range values of that particular grocery item.

Quality ratings were then generated for each option, in decreasing order, such that:

$$Q_i = Q_{Avg} + (Q_{Step} * \left(\left\lceil \frac{N}{2} \right\rceil - i\right))$$

where $Q_i$ is the price of item $i$, and $i$ ranges from 1 (the highest quality item) to $N$ (the lowest quality item), and where $N$ is the number of options in the choice set. This makes it so that the price of each item is equally spaced from one another and that the most extreme items have price values equivalent to the upper and lower range values of that particular grocery item.

For choice sets with three equi-preference contours, attributes for the most valuable contour were calculated the same as for choice sets with a single equi-preference contour, except that $N$ additionally divided by 3 as only a third of the options fall on this contour. Attributes for the least valuable contour are calculated as follows. A price step size was created to be equal to half the price step from the first, most valuable contour. Prices were then generated for each option, in decreasing order, such that:

$$P_i = \frac{P_{High} + P_{Avg}}{2} + (P_{Step} * \left(\left\lceil\frac{N}{2}\right\rceil - i\right))$$

where $P_i$ is the price of item $i$, and $i$ ranges from 1 (the most expensive item) to $N$ (the least expensive item), and where $N$ is the number of options in the contour. To generate quality ratings, a quality step was created to be equal to half the quality step from the first, more valuable contour.

Quality ratings were then generated for each option, in decreasing order, such that:

$$Q_i = \frac{Q_{High} + Q_{Avg}}{2} + (Q_{Step} * \left(\left\lceil\frac{N}{2}\right\rceil - i\right))$$

where $Q_i$ is the price of item $i$, and $i$ ranges from 1 (the highest quality item) to $N$ (the least expensive item), and where $N$ is the number of options in the contour.

The third, middle-valued contour was created by averaging the attribute values of the other two contours. Items were matched and averaged in pairs based on their rank order within each contour (e.g., the highest priced item from each), and their corresponding price and quality values were averaged to generate the attribute values for the middle contour.

For choice sets with randomized values, price and quality ratings for each option were generated using the RAND() function in excel and were designed to fall between the price and quality range values.

**An Integrated Neural Model of Value-based, Multiattribute Decision Making**

This dissertation investigated how the brain supports value-based decision making when individuals are confronted with choice sets that vary in size and structure. The empirical results highlighted distinct patterns of activation in the DLPFC, ACC, and AIns, suggesting that flexible decision strategies emerge from interactions among these regions. To contextualize these findings, I propose a neurocognitive model that builds on the sequential sampling framework of

Busemeyer et al. (2019) and the bottom-up, cue-driven model of strategy selection from Wichary and Smolen (2016). This model integrates these theoretical frameworks with neuroimaging data to explain how DLPFC, ACC, AIns, parietal cortex, and VMPFC contribute to multiattribute decision making. This model emphasizes the dynamic allocation of cognitive control and the modulation of valuation processes in response to context, including set size and the presence of salient alternatives.

The model is grounded in the following assumptions. Attributes are processed one at a time across alternatives through sequential sampling. Attribute values are integrated incrementally toward a decision threshold, with cognitive effort monitored throughout. Decision strategies shift dynamically in response to decision demands, detected conflict, or affective salience. The VMPFC maintains a continuous representation of option value based on accumulated evidence.

This model, like Busemeyer et al. (2019), assumes that attribute features of options are the inputs and that the parietal cortex processes these features. Previous research provides evidence for the parietal cortex in representing objective attributes (Wang et al., 2023).

Information about the choice set context is communicated from the parietal cortex to the ACC, which serves as a central hub of information integration from various brain regions. In our model, activity in the ACC serves as a signal for the amount of effort required to meet the demands of the decision process, essentially tracking difficulty. In the present study the ACC showed trending linear activity as a function of choice set size, because, even though the larger choice sets promoted a simplifying strategy, the cognitive resources required to implement the simpler strategy did not reduce the overall perceived difficulty of the choice process as reflected in the difficulty ratings of the post scan survey. In this model, the inverse U-shaped activity of the ACC in Reutskaja et al.'s (2018) choice reflected a much greater reduction in difficulty by the utilization of a simplifying decision strategy due to the nature of the aesthetic stimuli as well as the ability to rely on familiarity with the choice options. Participants in the Reutskaja et al. (2018) study had previously established their preferences for the distinct landscape images in a pre-scan rating task, allowing a simplifying strategy to much more readily reduce the difficulty of the scanner task. Indeed, for those participants who did rely on the value of dominance more in our study, we saw a reduction in the ACC activity, indicating that relying on dominance allowed for a reduction in difficulty for some of our participants.

The DLPFC in our model is responsible for actively maintaining and integrating attribute information in working memory. It implements top-down cognitive control, filtering irrelevant information and sustaining attention on task-relevant cues. In this way, a simplifying strategy used in large choice sets reduces the demand on the DLPFC because it reduces the number of options being considered and integrated. During choice sets of moderate size, peak DLPFC activity is seen because it is being maximally recruited to integrate information using a compensatory, weighted-additive strategy. Under extremely high load, the brain abandons exhaustive integration.

The VMPFC performs a comparison process, converting objective attribute information into subjective value representations in a common currency (Levy & Glimcher, 2012). The present study did not investigate the role of the VMPFC in choice overload, so we adopt the conceptualization of its involvement in value-based multiattribute choice outlined in Busemeyer et al. (2019). The VMPFC compares between options and attributes, providing a valuation of the alternatives that the DLPFC integrates into evidence accumulation.

The AIns is a key component of the brain's salience network, adding emotional significance to the decision process. We integrate the AIns as a signaler of affective heuristic bias, essentially a gut-level alert that an option or attribute stands out. When a particular option is salient (such as when one option dominates another), the AIns generates an affective response (through an aversion to the dominated option) that can prompt a heuristic choice. Notably, the AIns effect was more prominent in individuals who did not rely on a lexicographic decision strategy. Those using a compensatory strategy likely registered the dominance relationship more often. Thus, in our model the AIns works with the ACC as part of the salience/conflict monitoring system. The ACC provides an analytic conflict signal, while the insula provides an intuitive signal.

For this type of integrative model, there is a need for further specification in order to allow for testing and validation of its components. The current description is meant to provide a general framework for assessing the results of the neuroimaging study and tie them in with the mechanisms of previously existing models. Future research should aim to design neuroimaging studies that utilize reaction time information in order to fully integrate information about brain activity with sequential sampling models.

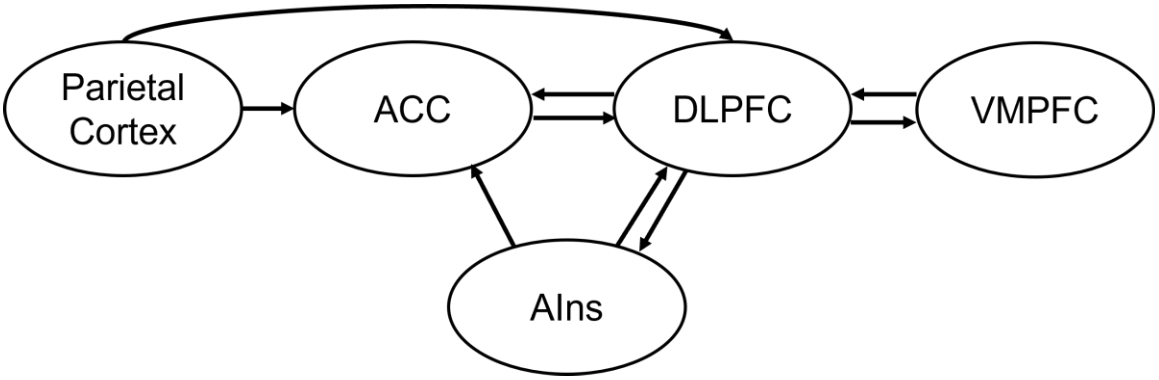


Supplemental Figure 1. Tentative Integrated Model Schematic. This schematic provides a visual representation of the tentative new integrative model. Arrows represent hypothesized functional interactions among key regions implicated in the present study and prior theoretical models. This schematic is intended as a conceptual framework as the model has not been formally tested.
